# A single BNT162b2 mRNA dose elicits antibodies with Fc-mediated effector functions and boost pre-existing humoral and T cell responses

**DOI:** 10.1101/2021.03.18.435972

**Authors:** Alexandra Tauzin, Manon Nayrac, Mehdi Benlarbi, Shang Yu Gong, Romain Gasser, Guillaume Beaudoin-Bussières, Nathalie Brassard, Annemarie Laumaea, Dani Vézina, Jérémie Prévost, Sai Priya Anand, Catherine Bourassa, Gabrielle Gendron-Lepage, Halima Medjahed, Guillaume Goyette, Julia Niessl, Olivier Tastet, Laurie Gokool, Chantal Morrisseau, Pascale Arlotto, Leonidas Stamatatos, Andrew T. McGuire, Catherine Larochelle, Pradeep Uchil, Maolin Lu, Walther Mothes, Gaston De Serres, Sandrine Moreira, Michel Roger, Jonathan Richard, Valérie Martel-Laferrière, Ralf Duerr, Cécile Tremblay, Daniel E. Kaufmann, Andrés Finzi

**Affiliations:** Centre de Recherche du CHUM, Montréal, QC, Canada; Département de Microbiologie, Infectiologie et Immunologie, Université de Montréal, Montreal, QC, Canada; Department of Microbiology and Immunology, McGill University, Montreal, QC, Canada; Fred Hutchinson Cancer Research Center, Vaccine and Infectious Disease Division, Seattle, WA, USA; University of Washington, Department of Global Health, Seattle, WA 98109, USA; Département des Neurosciences, Université de Montréal, Montreal, QC, Canada; Department of Microbial Pathogenesis, School of Medicine, New Haven, CT, USA; Institut National de Santé Publique du Québec, Quebec, QC, Canada; Laboratoire de Santé Publique du Québec, Institut national de santé publique du Québec, Sainte-Anne-de-Bellevue, QC, Canada; Department of Microbiology, New York University School of Medicine, New York, NY, USA; Département de Médecine, Université de Montréal, Montreal, QC, Canada; Consortium for HIV/AIDS Vaccine Development (CHAVD), La Jolla, CA, USA

**Keywords:** Coronavirus, COVID-19, SARS-CoV-2, Spike glycoproteins, mRNA vaccine, Variants, Antibodies, Humoral responses, Neutralization, ADCC, T-cell responses, Activation-induced marker assay, Intracellular cytokine staining

## Abstract

The standard dosing of the Pfizer/BioNTech BNT162b2 mRNA vaccine validated in clinical trials includes two doses administered three weeks apart. While the decision by some public health authorities to space the doses because of limiting supply has raised concerns about vaccine efficacy, data indicate that a single dose is up to 90% effective starting 14 days after its administration. We analyzed humoral and T cells responses three weeks after a single dose of this mRNA vaccine. Despite the proven efficacy of the vaccine at this time point, no neutralizing activity were elicited in SARS-CoV-2 naïve individuals. However, we detected strong anti-receptor binding domain (RBD) and Spike antibodies with Fc-mediated effector functions and cellular responses dominated by the CD4^+^ T cell component. A single dose of this mRNA vaccine to individuals previously infected by SARS-CoV-2 boosted all humoral and T cell responses measured, with strong correlations between T helper and antibody immunity. Neutralizing responses were increased in both potency and breadth, with distinctive capacity to neutralize emerging variant strains. Our results highlight the importance of vaccinating uninfected and previously-infected individuals and shed new light into the potential role of Fc-mediated effector functions and T cell responses in vaccine efficacy. They also provide support to spacing the doses of two-vaccine regimens to vaccinate a larger pool of the population in the context of vaccine scarcity against SARS-CoV-2.

## Introduction

The Severe Acute Respiratory Syndrome Coronavirus-2 (SARS-CoV-2) is the etiological agent of the Coronavirus disease 2019 (COVID-19), responsible for the current pandemic that infected over 120 million people and led to more than 2.66 million deaths worldwide^1, 2^. This pandemic caused a race for the elaboration of an effective vaccine against SARS-CoV-2^3, 4^. Currently approved vaccines target the highly immunogenic trimeric Spike (S) glycoprotein that facilitates SARS-CoV-2 entry into host cells via its receptor-binding domain (RBD) that interacts with angiotensin-converting enzyme 2 (ACE-2)^5, 6^. Among these vaccines, four are approved in many countries (Pfizer/BioNtech BNT162b2, Moderna mRNA-1273, AstraZeneca ChAdOx1 and Janssen Ad26.COV2S). The Pfizer/BioNtech BNT162b2 vaccine was developed using a novel technology based on mRNA^7^. This technology consists in intramuscular injection of a lipid nanoparticle-encapsulated synthetic mRNA vaccine encoding the viral Spike glycoproteins of SARS-CoV-2, which has shown to elicit a robust efficacy against the Wuhan-Hu-1 strain, which served as template for their development^8, 9^. This vaccine encodes for a membrane-anchored SARS-CoV-2 full-length spike, stabilized in a prefusion conformation by mutating the furin cleavage site and introducing two prolines in the S2 fusion machinery^7, 10^. However, the emergence of mutations in the SARS-CoV-2 S glycoprotein could affect different properties of the virus including affinity for its receptor, resulting in increased infectivity, transmissibility and evasion from humoral responses elicited by natural infection or vaccination^11^.

The D614G Spike mutation appeared very early in the pandemic and is now highly prevalent in all circulating strains^12^. The B.1.1.7 variant was first identified in the United Kingdom and has been spread rapidly to many countries since its identification. This variant contains several mutations in its S glycoproteins (ΔH69-V70, ΔY144, N501Y, A570D, P681H, T716I, S982A and D1118H) and has increased infectivity^13, 14^. Among the mutations present in the B.1.1.7 strain, the N501Y is also present in many other circulating variants (B.1.351 and P.1) and increases the affinity for the ACE-2 receptor^15, 16^. The E484K mutation, is part of the South African B.1.351 variant and is now found in several SARS-CoV-2 genomes worldwide that spread rapidly^17^. Studies have shown that this mutation increases affinity of the S glycoprotein for ACE-2^18^ and confers resistance to neutralization mediated by mAbs and plasma from naturally-infected and vaccinated individuals^19–22^. The S477N mutation confers a higher affinity for the ACE-2 receptor and has rapidly spread to many countries in Oceania and Europe^23–28^. The S477N and N501S mutations are found in several SARS-CoV-2 genomes in Quebec (Laboratoire de Santé Publique du Québec, unpublished data).

In spite of the proven clinical efficacy of BNT162b2, there are still limitations in the understanding of the protective components of the immune responses elicited by this vaccine. Such protection is mediated through a complex interplay between innate, humoral and cell-mediated immunity^29, 30^. Several reports showed that administration of the mRNA vaccine induced a strong humoral response after two doses, especially against the RBD domain^31, 32^. Robust CD4^+^ and CD8^+^ memory T cell responses are induced after SARS-CoV-2 infection^33, 34^ and play important roles in resolution of the infection^35^ including modulating disease severity in humans^36^ and reducing viral load in non-human primates (NHP)^37^. However, the detection of these specific memory T cells has been poorly studied in the SARS-CoV-2 vaccine development and represent a gap in the understanding of the induced cellular adaptative immune responses which are likely to also play an important role^8, 38^. Among CD4^+^ T cells, the T follicular helper (Tfh) subset is of particular interest, as it provides help for B cell maturation and development of high affinity antibody responses in the germinal center (GC) of secondary lymphoid organs. Studies have shown that a subset of CXCR5^+^ in blood, called circulating Tfh (cTfh)^39, 40^ has clonal, phenotypic and functional overlap with GC Tfh and reflect at least in part responses in tissues^41, 42^.

Results from phase III clinical trials have shown a vaccine efficacy of >90% starting 14 days after the injection of a single dose of BNT162b2 mRNA vaccine, thus before the administration of a second dose^7, 9, 43^. In this report, we characterized the humoral and T cell immune responses in cohorts of SARS-CoV-2 naïve and naturally-infected individuals prior and three weeks after a first dose of the BNT162b2 mRNA vaccine.

## Results

Here we analyzed humoral and cellular responses in blood samples from 16 SARS-CoV-2 naïve donors prior and after vaccination (median [range]: 21 days [16-26 days]). In addition, we examined the same immunological features in 16 individuals that were previously infected around 9 months before vaccination (median [range]: 266 days [116-309 days]) and three weeks after vaccination (median [range]: 21 days [17-25 days]). For 11 of these donors, we also longitudinally monitored evolution of the humoral response, from 6 weeks post-symptom onset (PSO, median [range]: 40 days [16-62 days]) to 3 weeks after vaccination. Basic demographic characteristics are summarized in Table 1. In the SARS-CoV-2 naïve group, the average age of donors was 48 years old (range: 21-59 years old), and samples were from 3 males and 13 females. In the group of previously-infected individuals, the average age of the donors was 48 years old (range: 23-65 years old), and samples were from 8 males and 8 females (Table 1).

**Table 1.** Characteristics of the vaccinated SARS-CoV-2 cohort.

| Group | n | Days between symptom onset and vaccination (median; day range) | Days after vaccination (median; day range) | Age (average; age range) | Sex |  |
| --- | --- | --- | --- | --- | --- | --- |
|  |  |  |  |  | F (n) | M (n) |
| **SARS-CoV-2 Naïve** | 16 | / | 21 (16 - 26) | 48 (21-59) | 13 | 3 |
| **SARS-CoV-2 Prior infection** | 16 | 266 (116-309) | 21 (17 - 25) | 48 (23-65) | 8 | 8 |

### Elicitation of SARS-CoV-2 antibodies against the full Spike and its receptor binding domain

To evaluate vaccine responses in SARS-CoV-2 naïve and previously-infected individuals, we first measured the presence of RBD-specific antibodies (IgG, IgM, IgA) using a previously described enzyme-linked immunosorbent (ELISA) RBD assay^44–46^. As expected, in the SARS-CoV-2 naïve group, we did not observe RBD-specific immunoglobulins (Ig) in samples recovered before vaccination (Figure 1A-D). Three weeks after the first dose, we found a significant increase in the total RBD-specific immunoglobulin levels with the exception of one donor from the SARS-CoV-2 naïve group who didn’t respond to the vaccine at this time-point. With the exception of IgM, vaccination induced similar levels of immunoglobulins (IgA and IgG) targeting the RBD to those present in individuals that were naturally infected 9 months ago (Figure 1A-D). In addition, RBD-specific IgM levels were significantly lower in the vaccinated SARS-CoV-2 naïve group compared to pre-vaccination samples from the previously-infected participants (Figure 1B).

**Figure 1.**
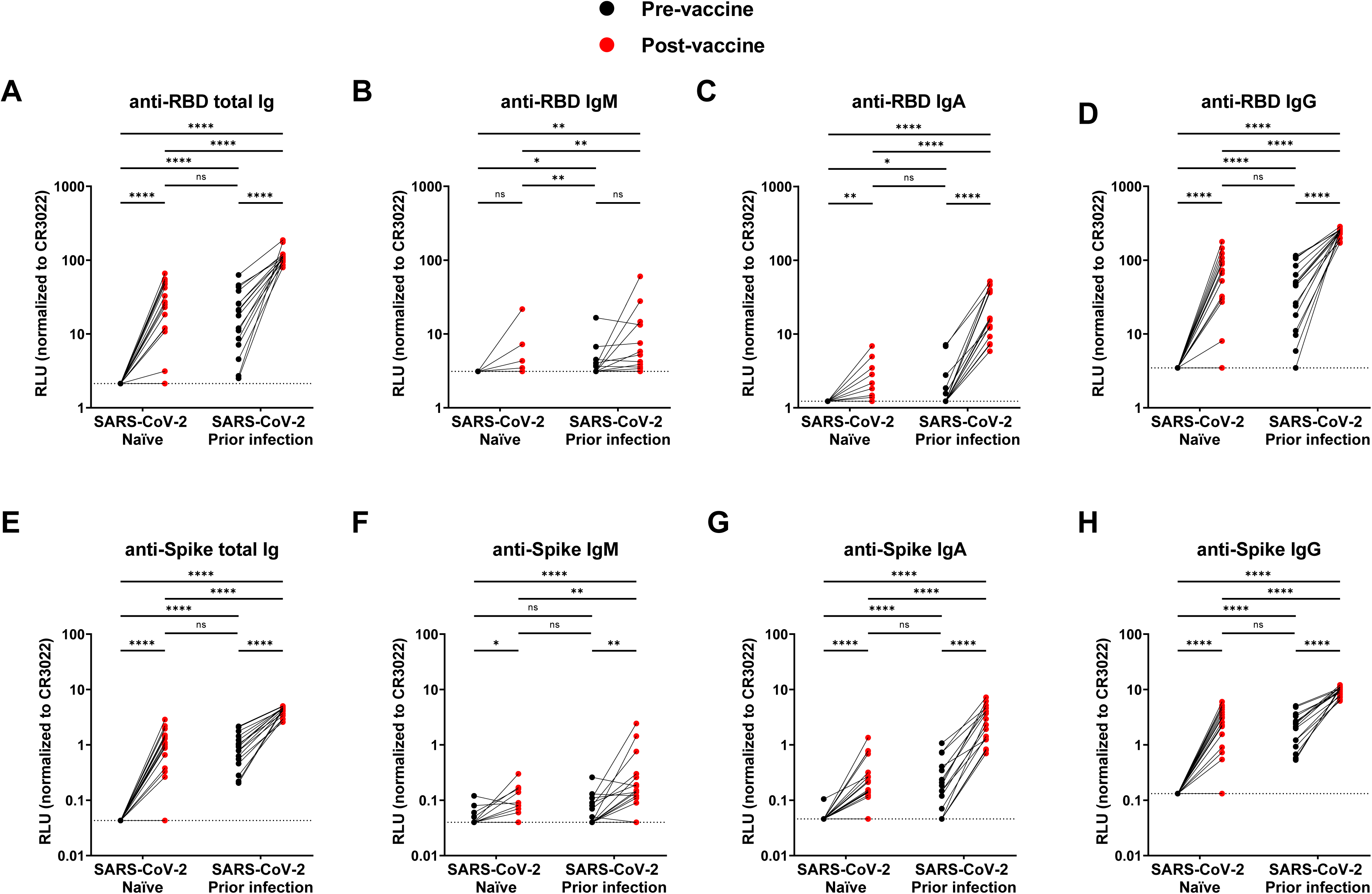
Elicitation of RBD- and Spike-specific antibodies by a single dose of Pfizer/BioNTech mRNA vaccine in SARS-CoV-2 naïve and previously-infected individuals. (**A-D**) Indirect ELISA was performed by incubating plasma samples from naïve and previously-infected donors collected before and after the first dose of vaccine with recombinant SARS-CoV-2 RBD protein. Anti-RBD antibody binding was detected using HRP-conjugated (**A**) anti-human IgM+IgG+IgA (**B**) anti-human IgM, (**C**) anti-human IgA, or (**D**) anti-human IgG. Relative light unit (RLU) values obtained with BSA (negative control) were subtracted and further normalized to the signal obtained with the anti-RBD CR3022 mAb present in each plate. (**E-H**) Cell-based ELISA was performed by incubating plasma samples from naïve and previously-infected donors collected before and after the first dose of vaccination with HOS cells expressing full-length SARS-CoV-2 Spike. Anti-Spike antibody binding was detected using HRP-conjugated (**E**) anti-human IgM+IgG+IgA (**F**) anti-human IgM, (**G**) anti-human IgA, or (**H**) anti-human IgG. RLU values obtained with parental HOS (negative control) were subtracted and further normalized to the signal obtained with the CR3022 mAb present in each plate. Limits of detection are plotted. (* P < 0.05; ** P < 0.01; *** P < 0.001; **** P < 0.0001; ns, non-significant).

In the group of individuals that were previously infected, despite a decline in the amount of Ig over time after infection (Figure S1A-D), most donors still had detectable anti-RBD-specific immunoglobulins just before vaccination, especially anti-RBD IgG (Figure 1A-D). For all participants, the first dose of vaccination led to a robust increase in anti-RBD IgG and anti-RBD IgA levels, higher than the first time point measured after the onset of symptoms (16-62 days; median: 40 days) (Figure 1C-D, S1C-D). Vaccination modestly increased the level of RBD-specific IgM (Figure 1B). Among the studied humoral responses, anti-SARS-CoV-2 neutralization returned to baseline most promptly, whereas ADCC remained more robust in the convalescent stage while still responding with a significant increase post vaccination (Figure S1E-G).

To evaluate if vaccine responses were limited to RBD or could be extended to antibodies against the full Spike glycoprotein, we used a cell-based ELISA (CBE) assay to measure levels of antibodies recognizing the native full-length S glycoprotein expressed at the cell surface^47^. In SARS-CoV-2 naïve individuals, the pattern was similar to that observed for the anti-RBD specific response, with a level of total Spike-specific immunoglobulins similar to that observed in previously-infected individuals before vaccination (Figure 1E). As we observed for anti-RBD Abs, vaccination of SARS-CoV-2 naïve individuals elicited higher levels of IgG than IgM or IgA (Figure 1F-H). The individual who did not elicit anti-RBD Abs upon vaccination didn’t elicit Abs against other regions of the Spike, with detection levels no higher than our seropositivity threshold level (Figure 1E-H).

Thus, vaccination in the SARS-CoV-2 naïve group elicited antibodies against the RBD and full Spike that reached similar levels than in naturally infected individuals 9 months post symptoms onset (Figure 1).

### Recognition of SARS-CoV-2 Spike variants and other *Betacoronaviruses*

SARS-CoV-2 is evolving, and variants of concern are emerging globally. Some harbor specific mutations in Spike that are associated with increased transmissibility and/or immune evasion^14, 48–51^. To evaluate whether a single dose of the Pfizer/BioNTech vaccine elicits antibodies that are able to recognize a broader spectrum of variants, including Spike with putative escape mutations, we measured the ability of plasma from vaccinated individuals to recognize different Spike variants expressed at the cell surface by flow cytometry, using a method we recently reported^44^. Briefly, 293T cells were transfected with plasmids expressing SARS-CoV-2 Spikes from the worldwide predominant strain (D614G)^52^, the B.1.1.7 variant or other individual concerning mutations (E484K, S477N, N501Y) , in parallel with Spike glycoproteins from other *Betacoronaviruses* (SARS-CoV-1, MERS-CoV, HCoV-OC43, HCoV-HKU1).Transfected cells were stained with plasma samples and incubated with secondary Abs recognizing all Ab isotypes. As expected, none of the SARS-CoV-2 naïve plasma samples obtained before vaccination (baseline) recognized the SARS-CoV-2 Spike (D614G) or any of its variants (Figure 2A-E). However, they were able to recognize Spikes from endemic human coronaviruses (HCoV-OC43, HCoV-HKU1) but not Spikes from highly pathogenic coronaviruses (SARS-CoV-1, MERS-CoV) (Figure 2F-I). In agreement with our CBE results (Figure 1), vaccination elicited antibodies that efficiently recognized the full Spike and all the tested variants (Figure 2A-E), except for the same donor that did not elicit RBD- or Spike-specific Abs. The recognition levels were equivalent to those observed for previously-infected individuals before vaccination. In the latter group, all plasma samples recognized the different Spike variants before vaccination, and the first dose of vaccine led to a significant increase in Spike recognition (Figure 2A-E). When we compared the differences in recognition between the SARS-CoV-2 variants, we observed that plasma from vaccinated SARS-CoV-2 naïve individuals recognized the different SARS-CoV-2 variants less efficiently compared to D614G Spike (Figure S2A). Plasma from previously infected individuals recognized all SARS-CoV-2 Spikes before and after vaccination. Vaccination however robustly enhanced recognition in this group, albeit a bit less efficiently for the Spike variants (Figure S2C).

**Figure 2.**
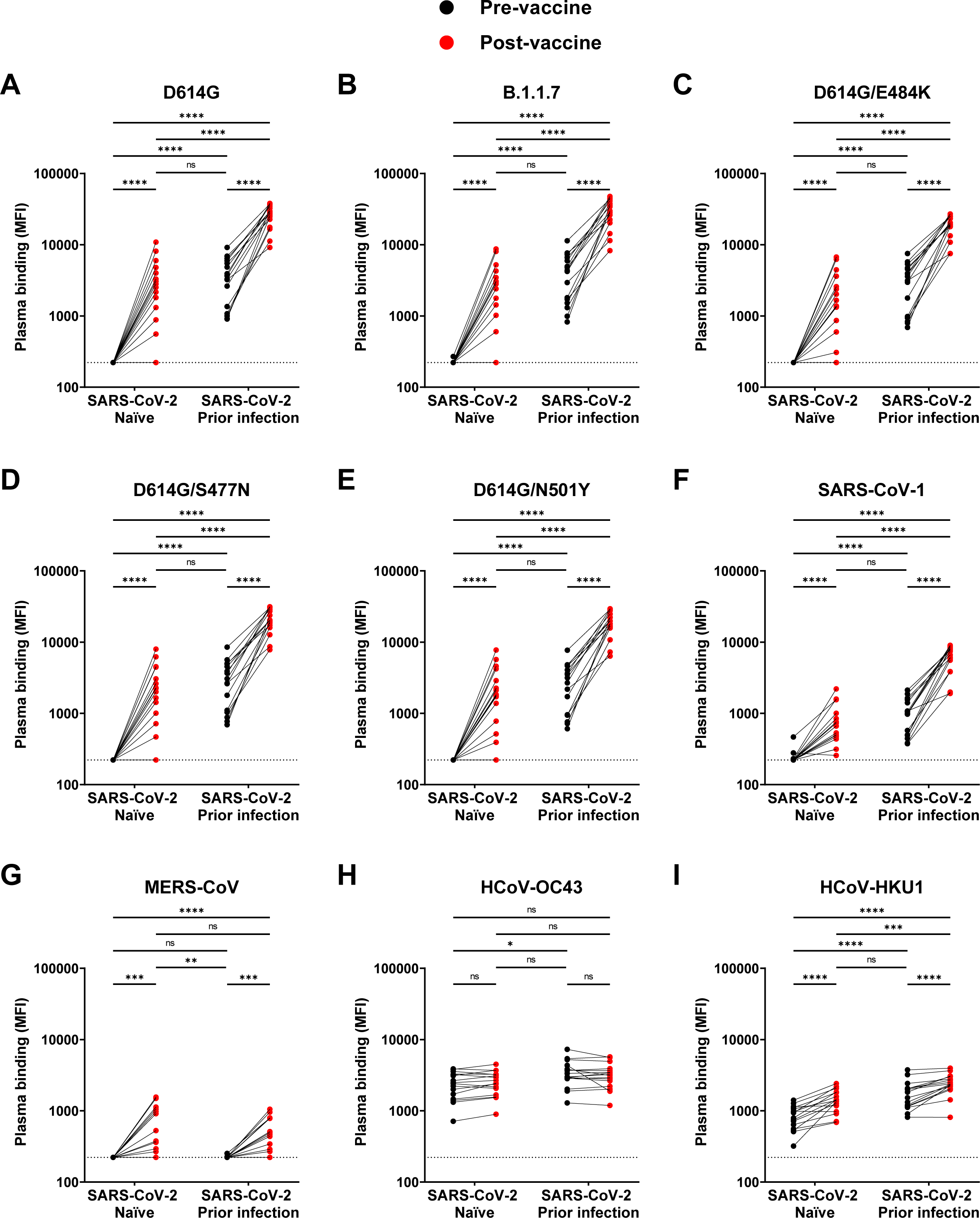
Detection of SARS-CoV-2 Spike variants and other *Betacoronaviruses*. (**A-I**) Cell-surface staining of 293T cells expressing full-length Spike from different SARS-CoV-2 variants and other human *Betacoronavirus* using plasma samples collected before and after first dose of vaccination in SARS-CoV-2 naïve and previously-infected donors. The graphs represent the median fluorescence intensities (MFI) obtained. Limits of detection are plotted. (* P < 0.05; ** P < 0.01; *** P < 0.001; **** P < 0.0001; ns, non-significant).

We recently reported that SARS-CoV-2 infection elicits cross-reactive antibodies that can recognize Spike from other human coronaviruses^44, 45^. To evaluate whether vaccination also elicited antibodies able to recognize Spike glycoproteins from other *Betacoronaviruses*, we evaluated the capacity of the different plasma samples to bind cell-surface expressed Spikes from SARS-CoV-1, MERS-CoV, HCoV-OC43 and HCoV-HKU1. As shown in Figure 2 (panels F-I), vaccination elicited cross-reactive antibodies in both groups with enhanced recognition of SARS-CoV-1, MERS-CoV and HCoV-HKU1 but not HCoV-OC43.

### Functional activities of vaccine-elicited antibodies

A single dose of the Pfizer/BioNTech vaccine was shown to be up to 90% efficacious starting two weeks after administration^7, 43, 53^. Among the immune responses elicited by the different SARS-CoV-2 vaccines, the neutralizing response is thought to be associated with vaccine efficacy^7, 54, 55^. To evaluate whether neutralizing responses were elicited within the first three weeks upon vaccine administration, we measured the capacity of plasma samples to neutralize pseudoviral particles carrying the SARS-CoV-2 Spike protein. Briefly, we incubated several dilutions of plasma with pseudoviruses before adding to 293T target cells stably expressing the human ACE-2 receptor, as we reported ^44–46, 56^. All pseudoviruses variants were infectious in this system with SARS-CoV-2 variants, particularly B.1.1.7 exhibiting enhanced infectivity (Figure S2D). We observed no neutralizing activity in plasma from vaccinated SARS-CoV-2 naïve individuals (Figure 3A), in agreement with previous findings^38^. As recently described^22, 57^, we observed that pre-existing SARS-CoV-2 neutralizing antibody responses were significantly boosted by a single dose of Spike-encoded mRNA vaccine (Figure 3A). Interestingly, a single dose enlarged the potency of the neutralizing response that was now able to efficiently neutralize pseudoviral particles bearing the B.1.1.7 Spike or from other variants with different concerning mutations (E484K, S477N, N501Y, N501S) (Figure S2E-G). Remarkably it also boosted neutralization activity against pseudoparticles bearing the SARS-CoV-1 Spike (Figure S2H).

**Figure 3.**
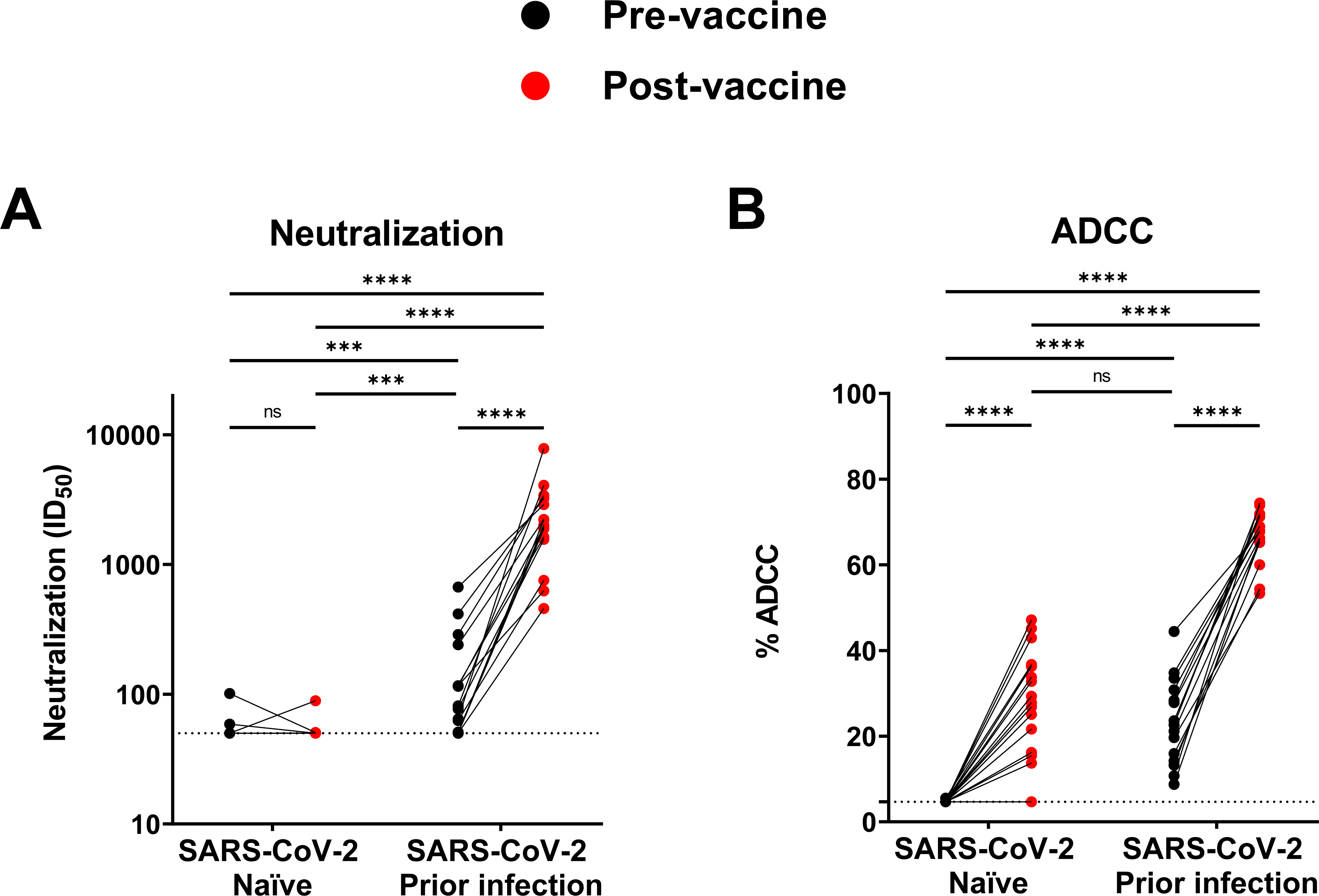
Neutralization and Fc-effector function activities in SARS-CoV-2 naïve and previously-infected individuals before and after a single dose of Pfizer/BioNTech mRNA vaccine. (**A**) Neutralizing activity was measured by incubating pseudoviruses bearing SARS-CoV-2 Spike glycoproteins, with serial dilutions of plasma for 1 h at 37°C before infecting 293T-ACE2 cells. Neutralization half maximal inhibitory serum dilution (ID_50_) values were determined using a normalized non-linear regression using GraphPad Prism software. (**B**) CEM.NKr parental cells were mixed at a 1:1 ratio with CEM.NKr-Spike cells and were used as target cells. PBMCs from uninfected donors were used as effector cells in a FACS-based ADCC assay. Limits of detection are plotted. (*** P < 0.001; **** P < 0.0001; ns, non-significant).

Since no neutralizing activity was detected in SARS-CoV-2 naïve vaccinated individuals, we decided to measure Fc-mediated effector functions that were also shown to play an important role against SARS-CoV-2 infection^58–60^. We used a human T-lymphoid cell line resistant to NK cell-mediated cell lysis (CEM.NKr) and stably expressing the full-length S protein on the cell surface (CEM.NKr-Spike) to measure antibody-dependent cellular cytotoxicity (ADCC). PBMCs from healthy individuals were used as effector cells. ADCC activity was measured by the loss of Spike-expressing GFP+ target cells, as we reported^47^. In agreement with the lack of Spike-specific antibodies, SARS-CoV-2 naïve individuals did not have detectable ADCC activity prior to vaccination (Figure 3B). The first dose of the vaccine induced a significant increase in ADCC activity, except for one sample, corresponding to the donor who had not developed anti-Spike antibodies. We noted that ADCC activity in vaccinated SARS-CoV-2 naïve individuals achieved comparable levels to those of previously-infected individuals before vaccination. Vaccination of this group significantly boosted the ADCC activity (Figure 3B). Based on these results it is tempting to speculate that the generation of antibodies with Fc-mediated effector functions, but without neutralization, might be sufficient to provide a certain level of protection.

### Spike-specific T cell vaccine responses differ between SARS-CoV-2 naïve and previously infected individuals

We examined whether prior SARS-CoV-2 infection alters the CD4^+^ and CD8^+^ T cell responses to vaccination. To measure SARS-CoV-2-Spike-specific T cells in the two cohorts of naïve persons and individuals with prior infection, we utilized two complementary methodologies, T cell receptor (TCR) dependent activation induced marker (AIM) assays and intracellular cytokine staining (ICS). We performed the cytokine-independent AIM assays as previously described^61^ with some modifications. We stimulated PBMC for 15h with an overlapping peptide pool spanning the Spike coding sequence and measured upregulation of the markers CD69, CD40L, 4-1BB and OX-40 upon stimulation. We used an AND/OR Boolean combination gating strategy to identify antigen-specific T cell responses (Figure S3A)^62^. We examined three populations of SARS-CoV-2-Spike-specific AIM^+^ T cells: (i) AIM^+^ total CD4^+^ T cells (Figure 4A), (ii) AIM^+^ circulating memory Tfh (cTfh) cells (Figure 4B) and (iii) AIM^+^ total CD8^+^ T cells (Figure 4C). We and others have shown that AIM assays can sensitively detect infection- and vaccine-induced cTfh responses^63, 64^, including in SARS CoV-2 infection^34^.

**Figure 4.**
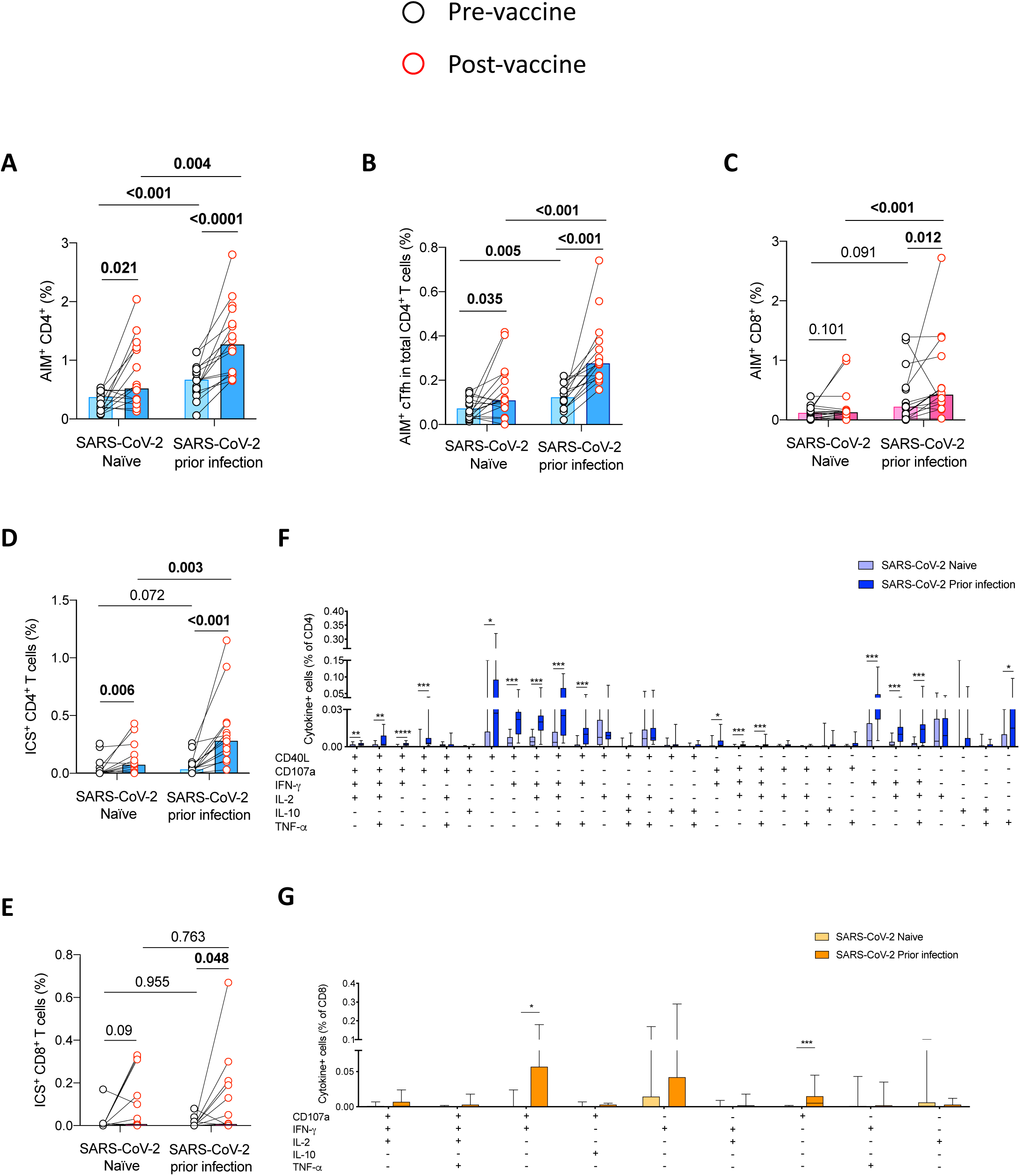
Spike-specific CD4^+^ and CD8^+^ T cell vaccine responses quantitatively and qualitatively differ in SARS-CoV-2 naïve versus previously-infected individuals. Net frequencies after Spike peptide pool stimulation of (**A**) total Spike-specific AIM^+^ CD4^+^ T cells, Spike-specific AIM^+^ cTfh (**C**) Spike-specific AIM^+^ CD8^+^ T cells in each donor prior to (V0) and post (V1) vaccination in the SARS-CoV-2 naïve participants and those with previous SARS-CoV-2 infection. Net frequencies of total S-specific responses measured by ICS for (**D**) CD4^+^ and (**E**) CD8^+^ T cells for each donor prior to and post vaccination. ICS^+^ populations include cells that expressed at least one cytokine and effector function upon 6h S peptide pool stimulation (CD40L, CD107a, IFN-γ, IL-2, IL-10 and TNF-α for CD4^+^; CD107a, IFN-γ, IL-2, IL-10 and TNF-α for CD8^+^ T cells). In (**A-E**), net frequency of the Spike-stimulated condition was calculated by subtracting the frequency detected in a DMSO control; bars correspond to median values and symbols represent biologically independent samples from n=16 SARS-CoV-2 naïve individuals and n=16 SARS-CoV-2 individuals with prior infection, lines connect data from the same donor. Analysis of the polyfunctionality of Spike-specific (**F**) CD4^+^ and (**G**) CD8^+^ T cells measured by ICS at the post vaccination (V1) time point. Data were analyzed by combinatorial gates based on the coexpression of CD40L, CD107a, IFN-γ, IL-2, IL-10 and TNF-α for CD4^+^ and CD107a, IFN-γ, IL-2, IL-10 and TNF-α for CD8^+^ T cells. Box-and-whisker plots show median values (line), 25^th^ to 75^th^ percentiles (box outline) and minimum and maximum values (whiskers). In (**F**) and (**G**) net frequency responses greater than 2-fold over DMSO control (background) were considered, significant *p*-values were indicated by * (* for <0.05; ** for <0.01, *** for <0.001). (**A**-**E**) P values were calculated by paired two-tailed Wilcoxon test for comparisons between the V0 and V1 time points in the same individual and Mann-Whitney for comparisons between the two cohorts at either the V0 or the V1 time point. (**F**-**G**) Comparisons between the polyfunctionality patterns were calculated using Mann-Whitney test.

After vaccination, we observed a significant increase in total Spike-specific AIM^+^CD4^+^ T cell responses in both groups of participants (Figure 4A) with significantly stronger responses in the prior infection group compared to the naïve group. We observed similar patterns with Spike-specific AIM^+^ cTfh responses which significantly increased after vaccination in both groups (Figure 4B) and stronger in the prior infection group compared to the naïve group. In contrast, there was a significant gain in Spike-specific AIM^+^ CD8^+^ T cell responses after vaccination only in the previous infected group. At the post-vaccination time points, AIM^+^ CD8^+^ T cell responses were also significantly higher than in the naïve group (Figure 4C). However, the frequencies range of these responses remains significantly lower than that of AIM^+^ CD4^+^ T cell responses regardless of the time point studied (Figure S3C).

To assess functionality and polarization of the SARS-CoV-2-Spike-specific T cell responses, we measured by ICS the cytokines secreted by CD4^+^ and CD8^+^ T cells in response to a 6h stimulation of PBMC with a Spike peptide pool. T cells were analyzed for expression of CD40L, CD107a, interferon (IFN)-γ, Interleukin (IL)-2, IL-10, IL-17A and tumor necrosis factor (TNF)-α. IL-17A expression was undetectable for most participants in both CD4^+^ and CD8^+^ T cell subsets, and CD40L negligible in CD8^+^ T cells. These subsets were thus not pursued, whereas all other functions were included in further analysis. We defined frequencies of cytokine^+^ CD4^+^ and CD8^+^ T cells as percentage of cells positive for one or more cytokines or functional markers (Figure S3B). Consistent with previous results on T cell responses specific for other viruses after natural infection^61, 63^ the magnitude of ICS^+^ T cells was lower than that of AIM^+^ T cell responses (Figure S3D-E), but there was a good correlation between both assays (Figure S3F). After vaccination, the ICS^+^ CD4^+^ and ICS^+^ CD8^+^ T cell responses were significantly increased only in the prior infection group (Figure 4D-E) with stronger responses compared to the naïve group. There were only trends for an increase in CD4^+^ and CD8^+^ T cell responses after this single dose of vaccine in the naïve cohort.

To qualitatively assess Spike-specific T cells in naïve and prior infection groups for polyfunctional responses after vaccination, we performed coexpression analysis using Boolean gating and examined each combinations of function (Figure 4F and G). In comparison to naïve individuals, dominant Spike-specific CD4^+^ T subsets that were preferentially increased by vaccination in prior infection included Spike-specific CD4^+^ T cells coexpressing CD40L, IFN-γ and TNF-α alone or in combination with other functions (CD107a, IL-2). The frequency of Spike-specific CD8^+^ T cells expressing IFN-γ or CD107a alone or combined together was also increased in prior infection compared to naïve participants.

These data show that while a single dose of mRNA vaccine could induce detectable Spike-specific CD4^+^ and CD8^+^ T cells in most individuals, including Spike-specific cTfh cells, independently of prior SARS-CoV-2 infection status, immunization elicited more robust and functionally skewed responses in participants with a history of SARS-CoV-2 infection, compared to naïve people, with preferential expansion of specific functional subsets.

### Relationship between SARS-CoV-2-Spike-specific T cell responses and humoral responses

Most protective antibody responses are dependent on CD4^+^ T cell help, which is critical for B cell expansion, affinity maturation and isotype switching. Therefore, we assessed whether pre-existing SARS-CoV-2-Spike-specific CD4^+^ T cells and cTfh responses were predictive of higher antibody titers and antibody functions, as measured by neutralization capacity and ADCC after vaccination, irrespective of prior infection status (Figure 5A). We observed different patterns of correlations between AIM^+^ CD4^+^, AIM^+^ cTfh and ICS^+^ CD4^+^ T cell frequencies measured before vaccination and vaccine-induced antibody responses (Figure 5A). We found that correlations between the function-agnostic AIM^+^ CD4^+^ T cell measurements and antibody responses were generally stronger than between ICS^+^ CD4^+^ T cell responses and serological measurements (Figure 5A). Notably, the Ig subsets measured after vaccination in the plasma of each participant showed significant positive correlations between pre-existing Spike-specific CD4^+^ T cell and cTfh responses on the one hand, and anti-Spike IgA and IgG post-vaccination on the other hand (Figure 5C, D, F and G). In contrast, we observed no significant correlations between total Spike-specific CD4^+^ T cell responses and anti-Spike IgM levels (Figure 5B) and between Spike-specific cTfh responses and anti-Spike IgM levels (Figure 5E). At the functional level, we observed significant correlations between all the pre-vaccination AIM^+^ Spike-specific memory CD4^+^ T cells and cTfh with ADCC and neutralization capacity post-immunization (Figure 5A).

**Figure 5.**
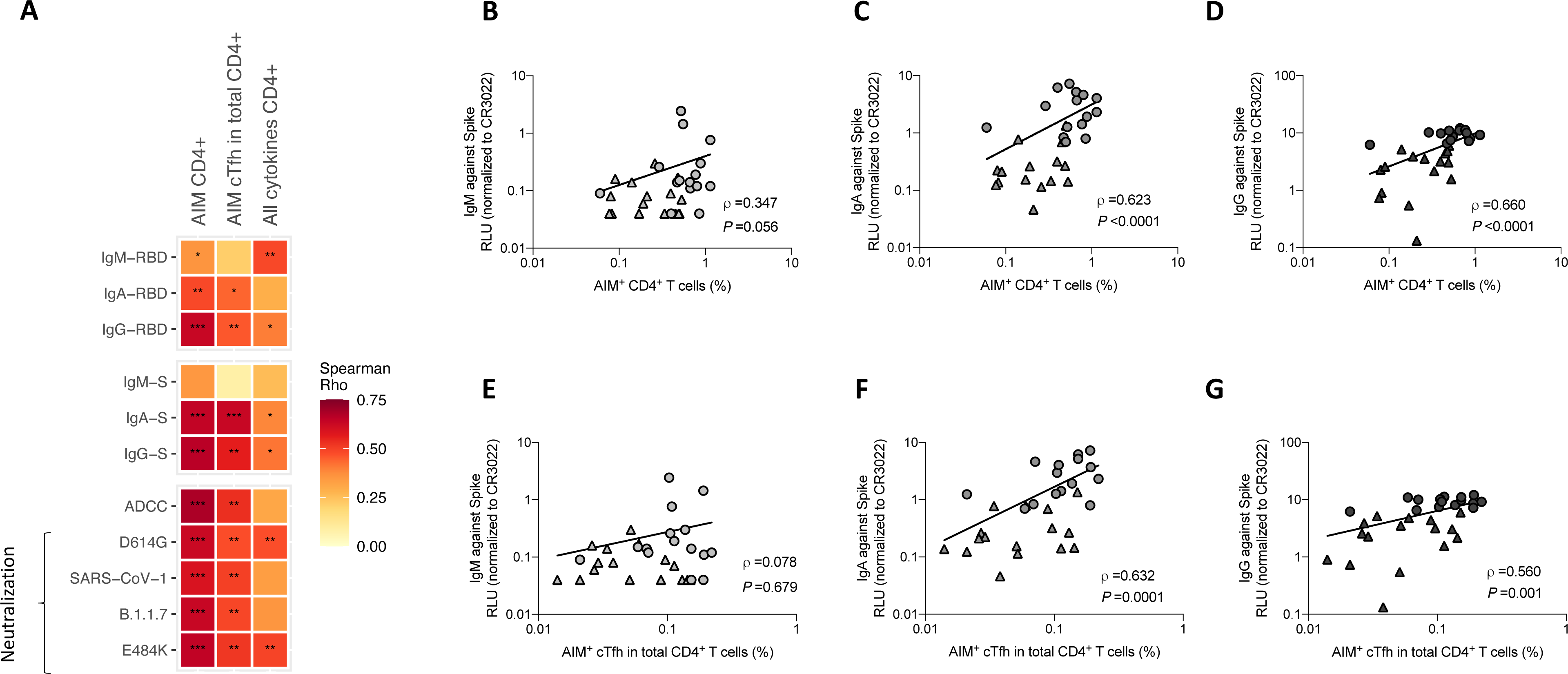
Total Spike-specific CD4^+^ T cells and Spike-specific cTfh responses at baseline correlate with humoral responses after vaccination. (**A**) Heatmap showing associations between total Spike-specific CD4^+^ T cell and Spike-specific cTfh responses at baseline (V0) and antibodies (against RBD and Spike), ADCC and neutralization functions after vaccination (V1). Color represents Rho value for each association calculated (Spearman correlation) and significant *p*-values were indicated by * (* for <0.05; ** for <0.01, *** for <0.001). Absence of significant correlations between IgM against Spike and AIM^+^ CD4^+^ T cells (**B**) and AIM^+^ cTfh responses (**E**). Positive correlations between IgA against Spike and AIM^+^ CD4^+^ T cells (**C**) and AIM^+^ cTfh responses (**F**). Positive correlation between IgG against Spike and AIM^+^ CD4^+^ T cells (**D**) and AIM^+^ cTfh responses (**G**). AIM^+^ cells were measured by flow cytometry and antibodies were quantified by CBE. Each symbol identifies one donor (SARS-CoV-2 naïve donors are represented by triangles and previously infected donors by circles).

These results suggest that pre-existing CD4^+^ T cell responses are beneficial for the generation of specific and effective humoral responses against SARS-CoV-2 after a single dose of mRNA vaccine, independently of prior SARS-CoV-2 infection.

### Evaluation of vaccine responses

Assessing the humoral responses revealed that the vaccine-induced responses (in naïve individuals) show striking similarities with the induced responses upon natural infection. With a few exceptions such as the neutralizing antibody response, at least for the given time points, the induced responses are similar (Figures 1-3). This translates into a similar network of pairwise correlations among all studied parameters when comparing discrete time points before vaccination in infected individuals and post vaccination in naïve individuals (Figure 6). As expected, naïve individuals before vaccination harbor hardly any interrelations between humoral and cellular anti-SARS-CoV-2 responses, which is in line with their overall low and unspecific absolute levels. Notably, when studying the effects of vaccination in previously infected individuals, the pairwise correlations are not getting stronger among our studied parameters, but the network of significant associations is broadened involving more interconnected parameters. It indicates that a heterogeneous boost, in this case a Spike mRNA vaccination boost upon natural infection as prime, brings in new flavors of host responses while diluting others.

**Figure 6.**
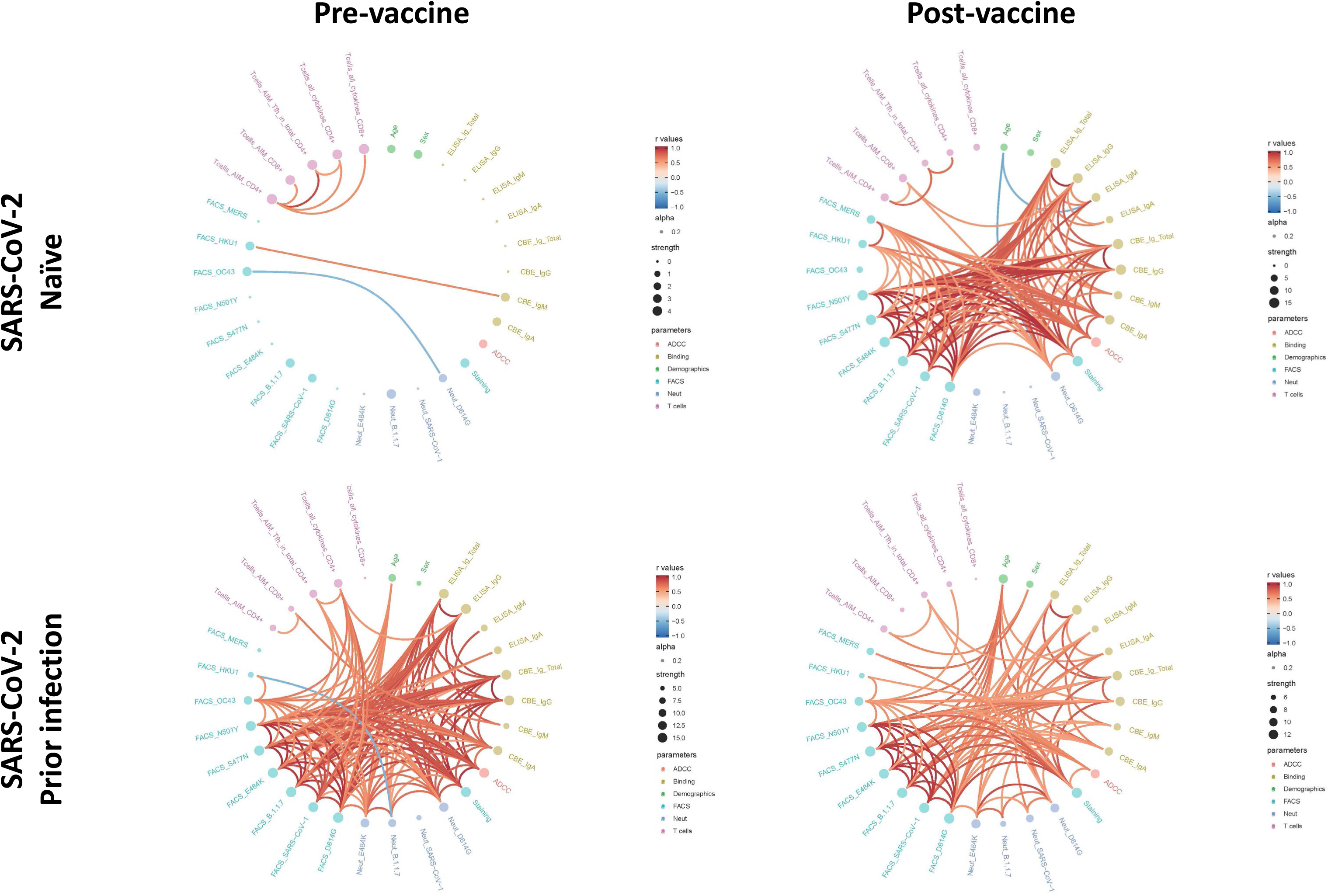
Mesh of correlations of humoral and cellular parameters at discrete time points before and after vaccination in SARS-CoV-2 naïve versus previously infected individuals. Edge bundling correlation plots where red and blue edges represent positive and negative correlations between connected parameters, respectively. Only significant correlations (p < 0.05, Spearman rank test) are displayed. Nodes are color-coded based on the grouping of parameters according to the legend. Node size corresponds to the degree of relatedness of correlations. Edge bundling plots are shown for correlation analyses using four different data sets, i.e., SARS-CoV-2 naïve and previously infected individuals before and after vaccination, respectively.

To investigate whether pre-existing humoral responses before vaccination predict the levels of induced/boosted responses upon vaccination, we performed a tandem correlation analysis focusing on pairs of correlations between time points before versus after vaccination (Figure S4). In naïve individuals, as expected, the low and SARS-CoV-2 unspecific responses before vaccination didn’t predict responses induced by vaccination. In contrast, individuals with previous SARS-CoV-2 infection harbor a much broader set of parameters pre-vaccination that predict induced responses post vaccination in the studied data set. Of interest, these correlations differ from the few observed in naïve individuals. In previously-infected individuals, most prominent patterns include the predictive value of binding, ADCC, and neutralization responses pre-vaccination for IgA responses in CBE assays and neutralization against viruses with the E484K Spike escape mutation post vaccination.

## Discussion

The mRNA vaccines have demonstrated a >90% efficacy starting 14 days after a single dose but the immune correlate of protection after a single dose remains unknown^7, 9, 43^. Here we measured several serological and cellular SARS-CoV-2-specific responses in SARS-CoV-2 naïve or previously-infected individuals. Surprisingly, despite the proven vaccine efficacy three weeks after vaccination^43, 53^ we observed no neutralizing activity in plasma from SARS-CoV-2 naïve vaccinated individuals. Neutralization is thought to play a central role in SARS-CoV-2 vaccine efficacy^7, 54, 55^, however, recent observations suggest that they might not be predictive, on their own, of protection^65^. Affinity maturation through germinal center selection can lead to more potent neutralizing antibody responses. While kinetics may differ according to the antigen used and route of administration, measurable neutralizing titers may take several weeks to develop in humans and NHP after immunization^66^, and even after neutralization titers begin to decrease, the somatic hypermutation (SHM) process can continue for months after acute SARS-CoV-2 infection^67^. Our results suggest that while the neutralization potency of vaccine-elicited antibodies is being developed, other antibody functions such as Fc-mediated effector functions could contribute to vaccine efficacy early on. Accordingly, three weeks after a single dose we observed strong ADCC but no neutralization activity (Figure 3). Strikingly, vaccination of previously-infected individuals induced a very significant increase of pre-existing humoral immunity including ADCC and neutralization. Neutralization potency was increased enabling neutralization of several variants including the B.1.1.7 variant, Spikes with the E484K mutation and even the phylogenetically more distant SARS-CoV-1.

We also demonstrated that the patterns of vaccine-induced T cell responses have analogies with those observed for antibody immunity. A single dose of BNT162b2 mRNA vaccine is also capable of generating SARS-CoV-2-specific T cell immune responses in both groups of individuals with a dominant CD4^+^ T cell responses, suggesting efficacy of the priming immunization in generating cellular immunity against SARS-CoV-2. However, we observed differences in the magnitude and quality of these responses between participants with and without prior infection. Individuals who had already encountered SARS-CoV-2 developed strong Spike-specific memory CD4^+^ and CD8^+^ T cells, consistent with secondary memory responses to a recall antigen. In contrast, naïve individuals showed significantly weaker Spike-specific CD4^+^ T cell responses and low to undetectable Spike-specific CD8^+^ T cell responses by AIM and ICS. Even though pre-vaccination T-cell responses to SARS-CoV-2 Spike glycoprotein were minimal in almost all naïve participants, it is still possible that the vaccine amplifies preexisting CD4^+^ cross-reactive T cell responses to endemic human coronaviruses. This suggests that a single dose of mRNA vaccine may be sufficient to elicit robust protective T cell responses in previously infected individuals, naïve persons will likely most benefit from repeat immunization.

Our results support the parallel use of both AIM and ICS assays for SARS-CoV-2 vaccine immunomonitoring. While most clinical trials relied on the IFN-γ ELISPOT assay and/or ICS to measure T cell responses, our data suggest the notion that BNT162b2 and some other SARS-CoV-2 vaccines in advanced clinical evaluation have demonstrated that vaccines for SARS-CoV-2 vaccines preferentially elicit Th1 responses may have to be reconsidered^68–71^. Indeed, these assays are sensitive for detection of Th1 cytokines and cytotoxic responses, but largely miss other important components of virus-specific cellular immunity. Consistent with this, we found that AIM assays detected vaccine-induced CD4^+^ and CD8^+^ T cells in natural infection^34^. Still, ICS assays were essential to reveal qualitative differences in cellular responses elicited after vaccination in previously infected versus naïve participants. With more proinflammatory and antiviral CD4^+^ and CD8^+^ T cell functional profiles in almost all previously-infected individuals, including IFN-γ, TNF-α, and for the CD8^+^ T cells, cytotoxic functions. Based on current knowledge, we suggest that a balanced humoral and Th1-directed cellular immune response may be important for protection from COVID-19 and the development of effective vaccine-induced immunization.

Spike-specific CD4^+^ T cell responses clearly dominated over CD8^+^ T cell responses, both for AIM and ICS measurements. Because of their role in antigen-specific B cell survival and maturation, we studied the correlation of CD4^+^ T cell responses with antibody immunity. We found strong positive correlations between Spike-specific AIM^+^ CD4^+^ T cell responses measured before vaccination and isotype-switched IgA and IgG antibody responses after vaccination, as well as ADCC and neutralization functions, contrasting with no significant correlations with the unswitched IgM responses. These patterns suggest that pre-existing memory T cell help is a major modulator of humoral SARS-CoV-2 vaccine responses. While the patterns of predictive associations were overall similar for total AIM^+^CD4^+^ T cells and AIM^+^ cTfh, the correlations were weaker with ICS measurements. Again, this suggests that the widely used ICS assays likely miss CD4^+^ T cell subsets that are important to sustain the development of vaccine antibody responses. Consistent with our observations on robust cTfh induction by BNT162b2 mRNA, it was shown that SARS-CoV-2 mRNA vaccine had a superior capacity, in comparison to rRBD-AddaVax, to elicit potent SARS-CoV-2-specific GC B cell responses after the administration of a single vaccine dose, suggesting that GC B cells and Tfh cells strongly correlated with the production of protective SARS-CoV-2-specific antibody responses^72^. Our results are also consistent with recent observations in convalescent COVID-19 donors, with reported correlations between antigen-specific CD4^+^ T cells^73^ and cTfh cells^74^ and SARS-CoV-2-specific antibodies. As the CD4 help-dependent development of high affinity antibody responses is a desired outcome after vaccination, our results provide clear rationales for assessing CD4^+^ T cell responses as part of the evaluation of SARS-CoV-2 vaccine immunogenicity and durability of protection, and for including function-agnostic techniques such as the AIM assays.

The availability of longitudinal sampling with six time points starting from a few weeks post symptoms onset up to three weeks post vaccination enabled us to investigate the predictive capacity of distinct time points in infected/convalescent individuals for vaccine outcome in terms of humoral responses (Figure S5). At the earliest time point, few weeks post symptoms onset, the predictive power of the studied parameters neutralization, ADCC, and ELISA binding responses (IgA, IgG, IgM, and total Ig) were low; however starting time point 2, total Ig, IgG, and ADCC responses gain power to significantly predict stronger IgG responses post vaccination. At the latest time point post symptoms onset, the predictive capacity of IgG and total Ig were partly diluted, but overall broadened, including predictions towards stronger IgA and IgM responses post vaccination.

We note that vaccination of SARS-CoV-2 naïve individuals bring their SARS-CoV-2 specific humoral and T cells responses to similar levels than the ones presented in individuals that were infected around nine months ago. Recent studies showed that natural infection confers up to 80% of protection from re-infection^75, 76^, however whether the same immune responses than those elicited by vaccination confer this protection remains unknown. These results give support to the consideration by various jurisdictions of a widened interval between the first and second dose in the context of vaccine shortage to protect a larger proportion of the population. The United Kingdom has decided to wait up to 12 weeks before administering the second dose of SARS-CoV-2 vaccines^77^ whereas Canada extended this interval up to 16 weeks^78^. This is also advocated in the United States in the context of the surging B.1.1.7 variant^79^. While the duration of a protective immune response elicited by a single dose of mRNA vaccines is unknown, given that memory is a core function of the immune system it is unlikely to decline within these intervals. Nevertheless, addressing this question will be very important as the larger the interval between doses the easier it will be to maximize the protection globally given the limited vaccine supply worldwide.

## Material and Methods

### Ethics Statement

All work was conducted in accordance with the Declaration of Helsinki in terms of informed consent and approval by an appropriate institutional board. Blood samples were obtained from donors who consented to participate in this research project at CHUM (19.381). Plasma and PBMCs were isolated by centrifugation and Ficoll gradient, and samples stored at -80°C and in liquid nitrogen, respectively, until use.

### Plasma and antibodies

Plasma from SARS-CoV-2 naïve and previously-infected donors were collected, heat-inactivated for 1 hour at 56°C and stored at -80°C until ready to use in subsequent experiments. Plasma from uninfected donors collected before the pandemic were used as negative controls and used to calculate the seropositivity threshold in our ELISA, cell-based ELISA, ADCC and flow cytometry assays (see below). The RBD-specific monoclonal antibody CR3022 was used as a positive control in our ELISA, cell-based ELISA, and flow cytometry assays and was previously described ^44, 45, 80^. Horseradish peroxidase (HRP)-conjugated antibodies able to detect all Ig isotypes (anti-human IgM+IgG+IgA; Jackson ImmunoResearch Laboratories) or specific for the Fc region of human IgG (Invitrogen), the Fc region of human IgM (Jackson ImmunoResearch Laboratories) or the Fc region of human IgA (Jackson ImmunoResearch Laboratories) were used as secondary antibodies to detect antibody binding in ELISA and cell-based ELISA experiments. Alexa Fluor-647-conjugated goat anti-human Abs able to detect all Ig isotypes (anti-human IgM+IgG+IgA; Jackson ImmunoResearch Laboratories) were used as secondary antibodies to detect plasma binding in flow cytometry experiments.

### Cell lines

293T human embryonic kidney cells (obtained from ATCC) were maintained at 37°C under 5% CO_2_ in Dulbecco’s modified Eagle’s medium (DMEM) (Wisent) containing 5% fetal bovine serum (FBS) (VWR) and 100 μg/ml of penicillin-streptomycin (Wisent). CEM.NKr CCR5+ cells (NIH AIDS reagent program) were maintained at 37°C under 5% CO_2_ in Roswell Park Memorial Institute (RPMI) 1640 medium (Gibco) containing 10% FBS and 100 μg/ml of penicillin-streptomycin. 293T-ACE2 and 293T-SARS-CoV-2 Spike cell lines were previously reported ^44^. HOS and CEM.NKr CCR5+ cells stably expressing the SARS-CoV-2 Spike glycoproteins were previously reported^47^.

### Protein expression and purification

FreeStyle 293F cells (Invitrogen) were grown in FreeStyle 293F medium (Invitrogen) to a density of 1 x 10^6^ cells/mL at 37°C with 8 % CO2 with regular agitation (150 rpm). Cells were transfected with a plasmid coding for SARS-CoV-2 S RBD using ExpiFectamine 293 transfection reagent, as directed by the manufacturer (Invitrogen). One week later, cells were pelleted and discarded. Supernatants were filtered using a 0.22 µm filter (Thermo Fisher Scientific). The recombinant RBD proteins were purified by nickel affinity columns, as directed by the manufacturer (Invitrogen). The RBD preparations were dialyzed against phosphate-buffered saline (PBS) and stored in aliquots at -80°C until further use. To assess purity, recombinant proteins were loaded on SDS-PAGE gels and stained with Coomassie Blue.

### Enzyme-Linked Immunosorbent Assay (ELISA)

The SARS-CoV-2 RBD ELISA assay used was previously described ^44, 45^. Briefly, recombinant SARS-CoV-2 S RBD proteins (2.5 μg/ml), or bovine serum albumin (BSA) (2.5 μg/ml) as a negative control, were prepared in PBS and were adsorbed to plates (MaxiSorp Nunc) overnight at 4°C. Coated wells were subsequently blocked with blocking buffer (Tris-buffered saline [TBS] containing 0.1% Tween20 and 2% BSA) for 1h at room temperature. Wells were then washed four times with washing buffer (Tris-buffered saline [TBS] containing 0.1% Tween20). CR3022 mAb (50ng/ml) or a 1/250 dilution of plasma from SARS-CoV-2-naïve or previously-infected donors were prepared in a diluted solution of blocking buffer (0.1 % BSA) and incubated with the RBD-coated wells for 90 minutes at room temperature. Plates were washed four times with washing buffer followed by incubation with secondary Abs (diluted in a diluted solution of blocking buffer (0.4% BSA)) for 1h at room temperature, followed by four washes. HRP enzyme activity was determined after the addition of a 1:1 mix of Western Lightning oxidizing and luminol reagents (Perkin Elmer Life Sciences). Light emission was measured with a LB942 TriStar luminometer (Berthold Technologies). Signal obtained with BSA was subtracted for each plasma and was then normalized to the signal obtained with CR3022 present in each plate. The seropositivity threshold was established using the following formula: mean of all SARS-CoV-2 negative plasma + (3 standard deviation of the mean of all SARS-CoV-2negative plasma).

### Cell-Based ELISA

Detection of the trimeric SARS-CoV-2 Spike at the surface of HOS cells was performed by a previously-described cell-based enzyme-linked immunosorbent assay (ELISA)^47^. Briefly, parental HOS cells or HOS-Spike cells were seeded in 96-well plates (4×10^4^ cells per well) overnight. Cells were blocked with blocking buffer (10 mg/ml nonfat dry milk, 1.8 mM CaCl_2_, 1 mM MgCl_2_, 25 mM Tris [pH 7.5], and 140 mM NaCl) for 30min. CR3022 mAb (1 μg/ml) or plasma from SARS-CoV-2 naïve or previously-infected donors (at a dilution of 1/250) were prepared in blocking buffer and incubated with the cells for 1h at room temperature. Respective HRP-conjugated antibodies were then incubated with the samples for 45 min at room temperature. For all conditions, cells were washed 6 times with blocking buffer and 6 times with washing buffer (1.8 mM CaCl_2_, 1 mM MgCl_2_, 25 mM Tris [pH 7.5], and 140 mM NaCl). HRP enzyme activity was determined after the addition of a 1:1 mix of Western Lightning oxidizing and luminol reagents (PerkinElmer Life Sciences). Light emission was measured with an LB942 TriStar luminometer (Berthold Technologies). Signal obtained with parental HOS was subtracted for each plasma and was then normalized to the signal obtained with CR3022 mAb present in each plate. The seropositivity threshold was established using the following formula: mean of all SARS-CoV-2 negative plasma + (3 standard deviation of the mean of all SARS-CoV-2negative plasma).

### Cell surface staining and flow cytometry analysis

293T cells were transfected with full length Spike of different *Betacoronavirus*. 48h post-transfection, S-expressing cells were stained with the CV3-25 Ab or plasma from SARS-CoV-2-naïve or previously-infected donors, prior and after vaccination (1/250 dilution). AlexaFluor-647-conjugated goat anti-human IgM+IgG+IgA Abs (1/800 dilution) were used as secondary antibodies. The percentage of transduced cells (GFP+ cells) was determined by gating the living cell population based on viability dye staining (Aqua Vivid, Invitrogen). Samples were acquired on a LSRII cytometer (BD Biosciences) and data analysis was performed using FlowJo v10.7.1 (Tree Star). The seropositivity threshold was established using the following formula: (mean of all SARS-CoV-2 negative plasma + (3 standard deviation of the mean of all SARS-CoV-2 negative plasma).

### ADCC assay

For evaluation of anti-SARS-CoV-2 antibody-dependent cellular cytotoxicity (ADCC), parental CEM.NKr CCR5+ cells were mixed at a 1:1 ratio with CEM.NKr.SARS-CoV-2.Spike cells. These cells were stained for viability (AquaVivid; Thermo Fisher Scientific, Waltham, MA, USA) and cellular dyes (cell proliferation dye eFluor670; Thermo Fisher Scientific) to be used as target cells. Overnight rested PBMCs were stained with another cellular marker (cell proliferation dye eFluor450; Thermo Fisher Scientific) and used as effector cells. Stained target and effector cells were mixed at a ratio of 1:10 in 96-well V-bottom plates. Plasma from SARS-CoV-2 naïve or previously-infected individuals (1/500 dilution) or monoclonal antibody CR3022 (1 µg/mL) were added to the appropriate wells. The plates were subsequently centrifuged for 1 min at 300xg, and incubated at 37°C, 5% CO2 for 5 hours before being fixed in a 2% PBS-formaldehyde solution. ADCC activity was calculated using the formula: [(% of GFP+ cells in Targets plus Effectors)-(% of GFP+ cells in Targets plus Effectors plus plasma/antibody)]/(% of GFP+ cells in Targets) x 100 by gating on transduced live target cells. All samples were acquired on an LSRII cytometer (BD Biosciences) and data analysis was performed using FlowJo v10.7.1 (Tree Star). The specificity threshold was established using the following formula: (mean of all SARS-CoV-2 negative plasma + (3 standard deviation of the mean of all SARS-CoV-2negative plasma).

### Plasmids

The plasmids expressing the human coronavirus Spikes of SARS-CoV-2, SARS-CoV-1^6, 81^, HCoV-OC43^44^ and MERS-CoV^82^ were previously reported. The HCoV-HKU1 Spike expressing plasmid was purchased from Sino Biological. SARS-CoV-2 Spike mutations were introduced using the QuikChange II XL site-directed mutagenesis protocol (Stratagene). The presence of the desired mutations was determined by automated DNA sequencing. The plasmid encoding the Spike of theB.1.1.7 variant was codon-optimized and synthesized by Genscript.

### Viral infectivity

293T cells were transfected with the lentiviral vector pNL4.3 R-E-Luc (NIH AIDS Reagent Program) and plasmid encoding for the indicated Spike glycoprotein (D614G, B.1.1.7, D614G/E484K, D614G/N501S, D614G/S477N and D614G/N501Y) at a ratio of 5:4. Two days post-transfection, cell supernatants were harvested and stored at -80°C until use. The RT activity was evaluated by measure of the incorporation of [*methyl*-^3^H]TTP into cDNA of a poly(rA) template in the presence of virion-associated RT and oligo(dT). Normalized amount of RT activity pseudoviral particles were added to 293T-ACE2 target cells for 48h at 37°C. Then, cells were lysed by the addition of 30 µL of passive lysis buffer (Promega) followed by one freeze-thaw cycle. An LB942 TriStar luminometer (Berthold Technologies) was used to measure the luciferase activity of each well after the addition of 100 mL of luciferin buffer (15mM MgSO_4_, 15mM KPO_4_ [pH 7.8], 1mM ATP, and 1mM dithiothreitol) and 50 mL of 1mM d-luciferin potassium salt (Thermo Fisher Scientific). RLU values obtained were normalized to D614G.

### Virus neutralization assay

293T cells were transfected with the lentiviral vector pNL4.3 R-E-Luc (NIH AIDS Reagent Program) and a plasmid encoding for the indicated Spike glycoprotein (D614G, B.1.1.7, D614G/E484K, D614G/N501S, D614G/S477N, D614G/N501Y and SARS-CoV-1) at a ratio of 5:4. Two days post-transfection, cell supernatants were harvested and stored at -80°C until use. 293T-ACE2 target cells were seeded at a density of 1×10^4^ cells/well in 96-well luminometer-compatible tissue culture plates (Perkin Elmer) 24h before infection. Pseudoviral particles were incubated with the indicated plasma dilutions (1/50; 1/250; 1/1250; 1/6250; 1/31250) for 1h at 37°C and were then added to the target cells followed by incubation for 48h at 37°C. Then, cells were lysed by the addition of 30 µL of passive lysis buffer (Promega) followed by one freeze-thaw cycle. An LB942 TriStar luminometer (Berthold Technologies) was used to measure the luciferase activity of each well after the addition of 100 mL of luciferin buffer (15mM MgSO_4_, 15mM KPO_4_ [pH 7.8], 1mM ATP, and 1mM dithiothreitol) and 50 mL of 1mM d-luciferin potassium salt (Thermo Fisher Scientific). The neutralization half-maximal inhibitory dilution (ID_50_) represents the plasma dilution to inhibit 50% of the infection of 293T-ACE2 cells by SARS-CoV-2 pseudoviruses.

### Intracellular Cytokine Staining

PBMCs were thawed and rested for 2 h in RPMI 1640 medium supplemented with 10% FBS, Penicillin-Streptomycin (Thermo Fisher scientific, Waltham, MA) and HEPES (Thermo Fisher scientific, Waltham, MA). 2×10^6^ PBMCs were stimulated with a Spike glycoprotein peptide pool (0.5 μg/ml per peptide from JPT, Berlin, Germany) corresponding to the pool of 315 overlapping peptides (15-mers) spanning the complete amino acid sequence of the Spike glycoprotein. Cell stimulation was carried out for 6h in the presence of mouse anti-human CD107A, Brefeldin A and monensin (BD Biosciences, San Jose, CA) at 37 °C and 5% CO_2_. DMSO-treated cells served as a negative control. Cells were stained for aquavivid viability marker (Thermo Fisher scientific, Waltham, MA) for 20 min at 4 °C and surface markers (30 min, 4 °C), followed by intracellular detection of cytokines using the IC Fixation/Permeabilization kit (Thermo Fisher scientific, Waltham, MA) according to the manufacturer’s protocol before acquisition on a Symphony flow cytometer (BD Biosciences, San Jose, CA). Antibodies used for surface and intracellular staining are listed in the Supplemental Table 2. Stained PBMCs were acquired on Symphony cytometer (BD Biosciences) and analyzed using FlowJo v10.7.1 software.

### Activation-induced marker assay

PBMCs were thawed and rested for 3h in 96-well flat-bottom plates in RPMI 1640 supplemented with HEPES, penicillin and streptomycin and 10% FBS. 1.7×10^6^ PBMCs were stimulated with a Spike glycoprotein peptide pool (0.5 μg/ml per peptide) for 15h at 37 °C and 5% CO_2_. A DMSO-treated condition served as a negative control and SEB-treated condition (0.5 μg/ml) as positive control. Cells were stained for viability dye for 20min at 4 °C then surface markers (30 min, 4 °C) (See Supplementary Table 3 for antibody staining panel). Cells were fixed using 2% paraformaldehyde before acquisition on Symphony cytometer (BD Biosciences). Analyses were performed using FlowJo v10.7.1 software.

### Statistical analysis

Symbols represent biologically independent samples from SARS-CoV-2 naïve individuals (n=16) and SARS-CoV-2 prior infection individuals (n=16). Lines connect data from the same donor. Statistics were analyzed using GraphPad Prism version 8.0.1 (GraphPad, San Diego, CA). Every dataset was tested for statistical normality and this information was used to apply the appropriate (parametric or nonparametric) statistical test. Differences in responses for the same patient before and after vaccination were performed using Wilcoxon matched pair tests. Differences in responses between naïve and previously-infected individuals were measured by Mann-Whitney tests. Differences in responses against the SARS-CoV-2 variants for the same patient were measured by Friedman test. P values < 0.05 were considered significant; significance values are indicated as ∗ p < 0.05, ∗∗ p < 0.01, ∗∗∗ p < 0.001, ∗∗∗∗ p < 0.0001. Line charts were created with Prism 8.4.3 using normalized data and Akima spline interpolation. For correlations, Spearman’s R correlation coefficient was applied. Statistical tests were two-sided and p<0.05 was considered significant.

### Software scripts and visualization

Normalized heatmaps were generated using the complexheatmap, tidyverse, and viridis packages in R and RStudio^83, 84^ . Normalizations were done per “Analysis group”, e.g., separately for all neutralization data, T cell responses, etc, except for binding analysis, which was normalized per individual parameter because different antibodies are needed for the detection of IgG, IgM and IgA responses. IDs were grouped and clustered separately according to naïve versus previously-infected individuals, and also according to the time points before vaccination (V0) and after vaccination (V1). Squared correlograms were generated using the corrplot and RColorBrewer packages in program R and RStudio. Edge bundling graphs were generated in undirected mode in R and RStudio using ggraph, igraph, tidyverse, and RColorBrewer packages. Edges are only shown if p < 0.05, and nodes are sized according to the connecting edges’ r values. Nodes are color-coded according to groups of parameters. Area graphs were generated for the display of normalized time series. The plots were created in RawGraphs using DensityDesign interpolation and vertically un-centered values^85^ . Line charts in overlay were created with Prism 8.4.3 using normalized data per response and Akima spline interpolation.

## Acknowledgements

The authors are grateful to the donors who participated in this study. The authors thank the CRCHUM BSL3 and Flow Cytometry Platforms for technical assistance. We thank Dr. Stefan Pöhlmann (Georg-August University, Germany) for the plasmid coding for SARS-CoV-2 and SARS-CoV-1 S glycoproteins and Dr. M. Gordon Joyce (U.S. MHRP) for the monoclonal antibody CR3022. This work was supported by le Ministère de l’Économie et de l’Innovation du Québec, Programme de soutien aux organismes de recherche et d’innovation to A.F. and by the Fondation du CHUM. This work was also supported by a CIHR foundation grant #352417 to A.F., by an Exceptional Fund COVID-19 from the Canada Foundation for Innovation (CFI) #41027 to A.F. and D.E.K. Work on variants presented here was supported by the Sentinelle COVID Quebec network led by the LSPQ in collaboration with Fonds de Recherche du Québec Santé (FRQS) to A.F and Genome Canada – Génome Québec to M.R. Support was also provided by NIH AI144462 CHAVD to D.E.K. (Consortium PI: Dennis Burton). A.F. is the recipient of Canada Research Chair on Retroviral Entry no. RCHS0235 950-232424. V.M.L. is supported by a FRQS Junior 1 salary award. D.E.K. is a FRQS Merit Research Scholar. R.D. was supported by NIH grant R01 AI122953-05. S.P.A, J.P. and G.B.B. are supported by CIHR fellowships. R.G. is supported by a MITACS Accélération postdoctoral fellowship. The funders had no role in study design, data collection and analysis, decision to publish, or preparation of the manuscript. We declare no competing interests.

## Supplemental Information

Supplemental information includes 5 figures and 3 tables and can be found online.

**Supplemental Figure 1.**
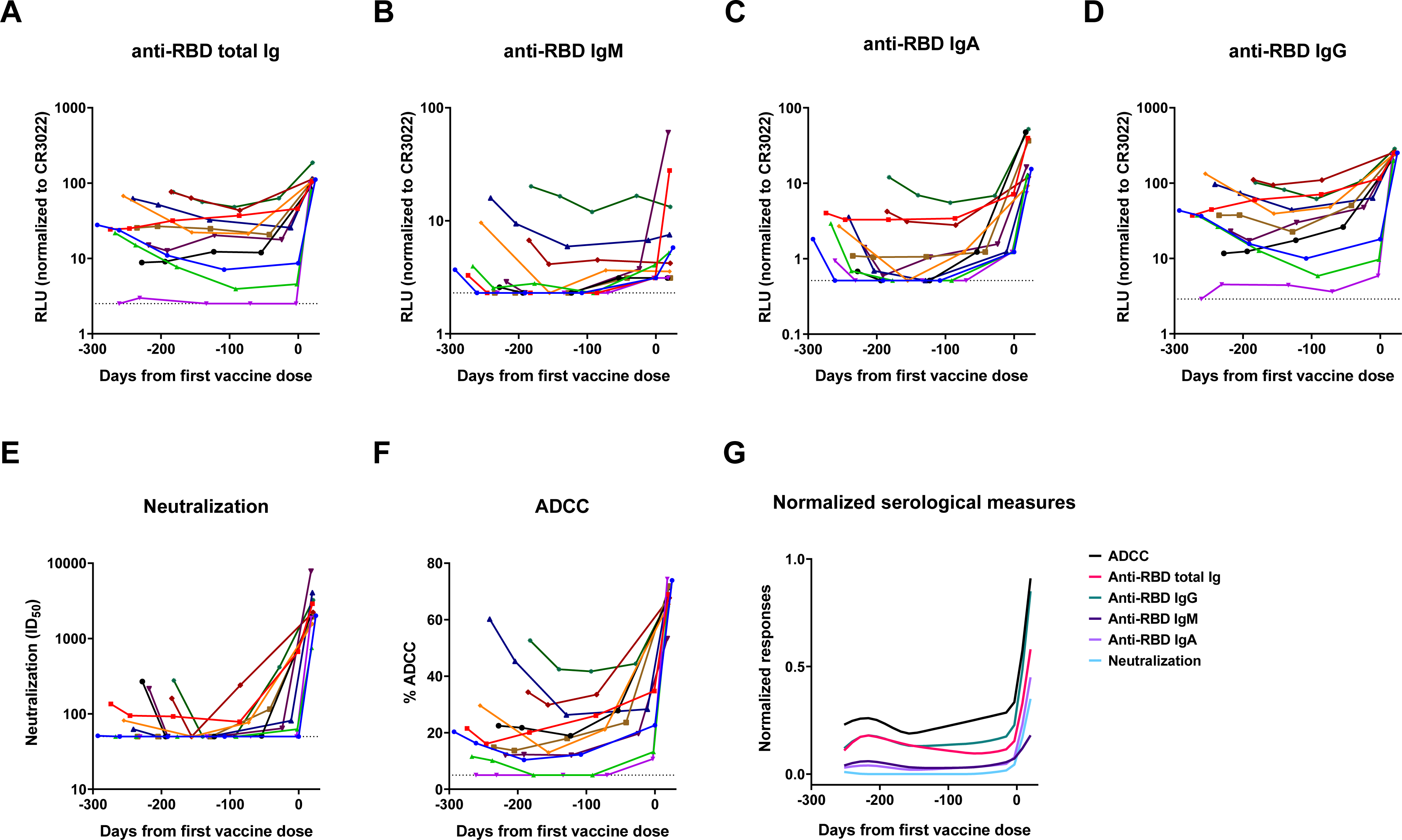
Longitudinal humoral responses in previously-infected SARS-CoV-2 individuals. Serological samples from eleven individuals that were previously infected were collected at different time points after symptoms onset (between 16- and 309-days post-symptoms onset) and three weeks after vaccination^37^. (**A-D**) RBD ELISA. Anti-RBD antibody binding was detected using HRP-conjugated (**A**) anti-human IgM+IgG+IgA (**B**) anti-human IgM, (**C**) anti-human IgA, or (**D**) anti-human IgG. Relative light unit (RLU) values obtained with BSA (negative control) were subtracted and further normalized to the signal obtained with the anti-RBD CR3022 present in each plate, as described in the material and methods section. (**E**) Neutralizing activity was measured by incubating pseudoviruses with serial dilutions of plasma for 1 h at 37°C before infecting 293T-ACE2 cells. Neutralization half maximal inhibitory serum dilution (ID50) values were determined using a normalized non-linear regression using GraphPad Prism software. (**F**) CEM.NKr parental cells were mixed at a 1:1 ratio with CEM.NKr-Spike cells and were used as target cells. PBMCs from uninfected donors were used as effector cells in a FACS-based ADCC assay. (**G**) Line charts showing normalized immune responses in overlay over the study period from 293 days before until 25 days post SARS-CoV-2 vaccination in individuals with prior SARS-CoV-2 infection. Curves were generated using Monotone X interpolation of data points. Time point of vaccination is displayed at X=0. Limits of detection are plotted.

**Supplemental Figure 2.**
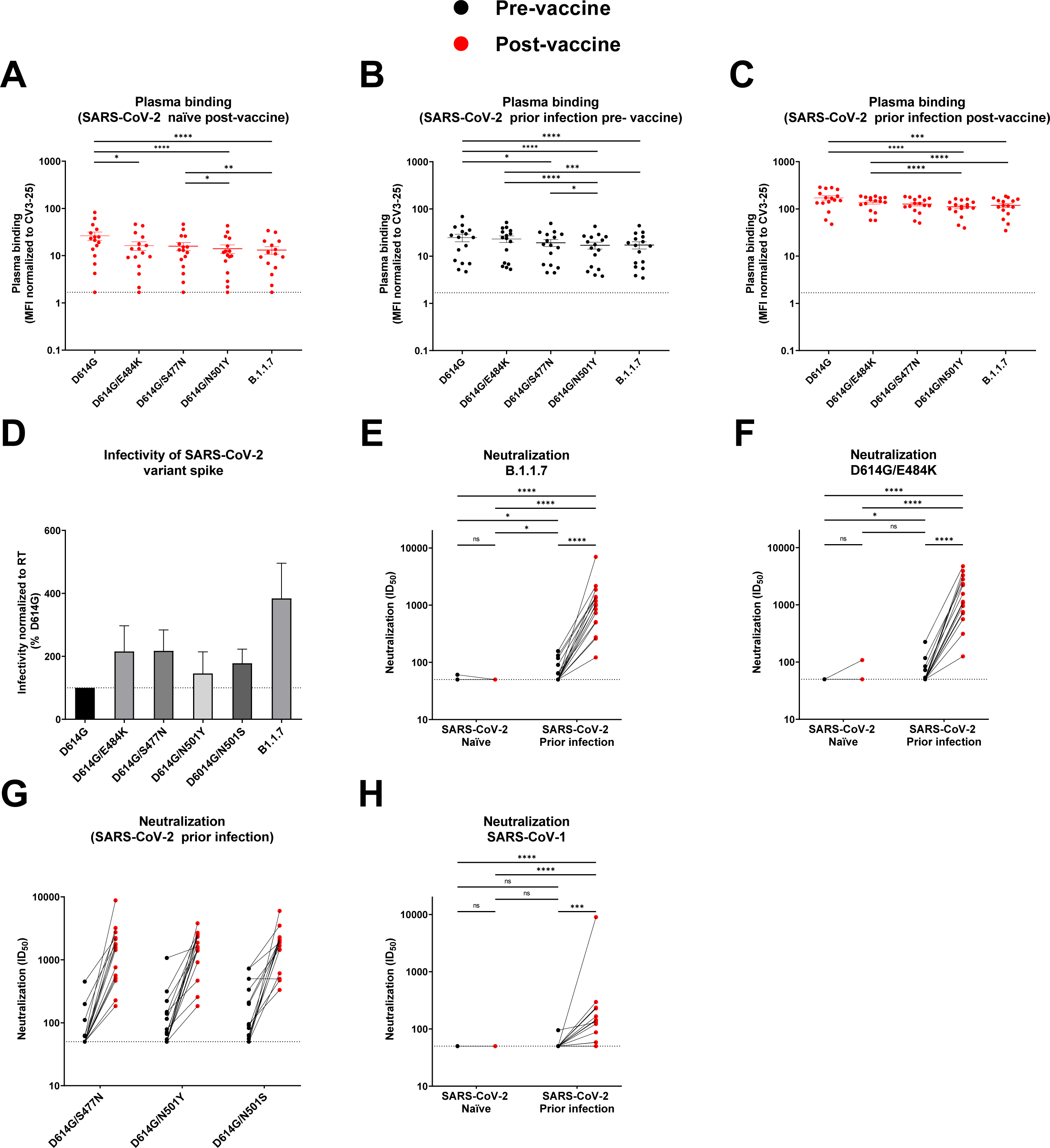
Impact of SARS-CoV-2 mutations on vaccine elicited humoral responses. (**A-C**) Cell-surface staining of 293T cells expressing full-length Spike from different SARS-CoV-2 variants using plasma samples collected in (**A**) SARS-CoV-2 naïve donors after first dose of vaccine, in previously-infected donors (**B**) before and (**C**) after vaccination. The graphs represent the median fluorescence intensities (MFI) obtained normalized to the MFI obtained with the CV3-25 Ab. (**D**) Pseudoviral particles bearing SARS-CoV-2 S glycoproteins from different variants were used to infect 293T-ACE2 cells for 2 days at 37°C. RLU values obtained were normalized to D614G. These experiments were repeated three times. Error bars indicate means ± SEM. (**E-H**) Neutralizing activity was measured by incubating indicated pseudoviruses with serial dilutions of plasma for 1 h at 37°C before infecting 293T-ACE2 cells. Neutralization half maximal inhibitory serum dilution (ID50) values were determined using a normalized non-linear regression using GraphPad Prism software. Limits of detection are plotted. (* P < 0.05; ** P < 0.01; *** P < 0.001; **** P < 0.0001; ns, non-significant).

**Supplemental Figure 3.**
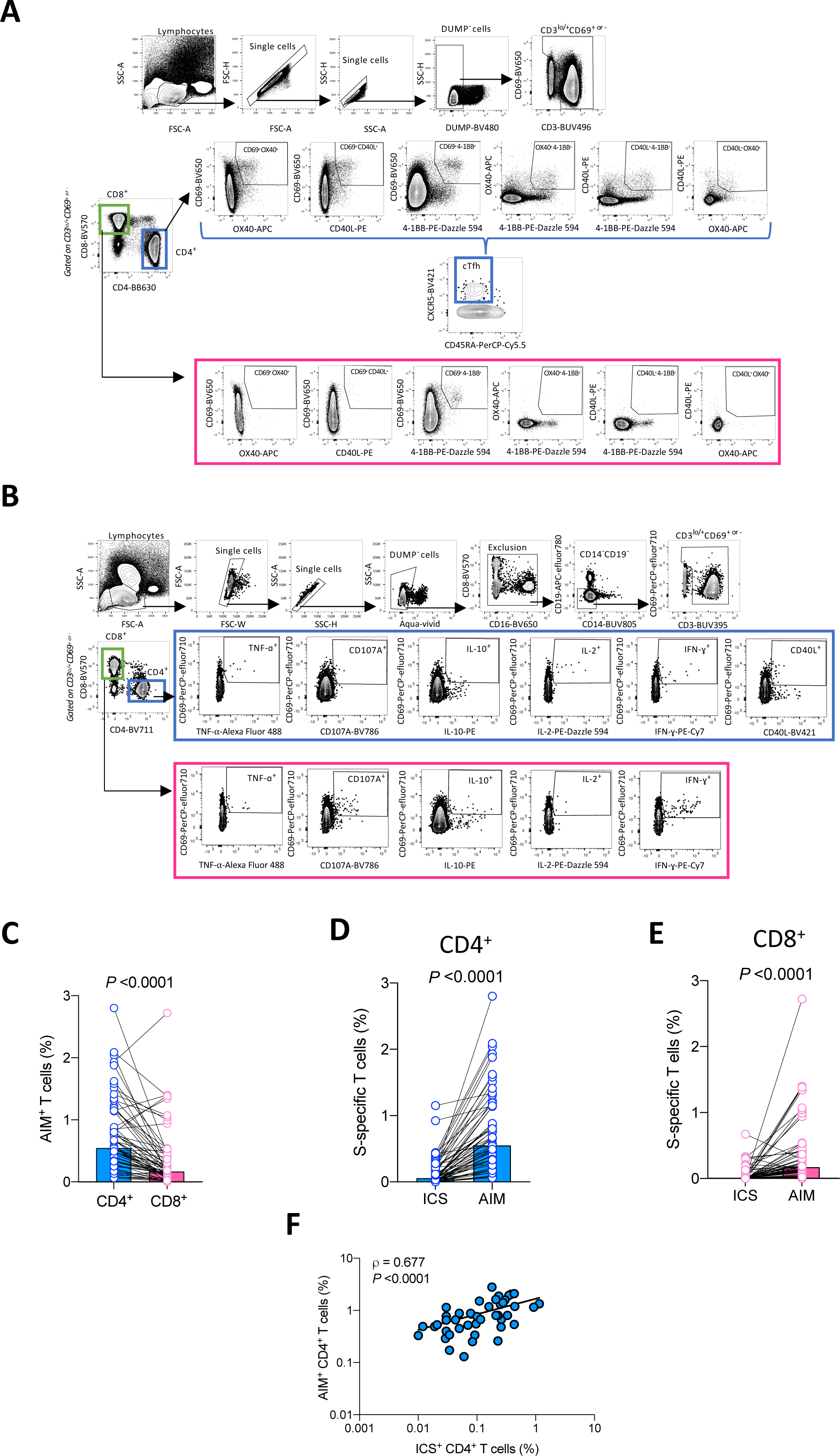
Gating strategy of measurements of Spike-specific T cell responses and comparisons of AIM and ICS assays. Representative flow cytometry gates to identify Spike-specific T cells in PBMCs from naïve and previously-infected donors. (**A**) Boolean OR gating strategy were used to analyze activation-induced markers (AIM^+^) Spike-specific responses in CD4^+^ T cells and cTfh (in blue) and CD8^+^ T cells (in pink) after a 15h stimulation with a Spike peptide pool. AIM^+^ T cells include cells that were CD69^+^OX40^+^ or CD69^+^CD40L^+^ or CD69^+^4-1BB^+^ or OX40^+^4-1BB^+^ or CD40L^+^4-1BB^+^ or CD40L^+^OX40^+^. (**B**) Boolean OR gating strategies were used to analyze by intracellular staining (ICS) the cytokine/effector functions in CD4^+^ (in blue) and CD8^+^ (in pink) T cells and identify T cells that responded to Spike peptide pool after 6h stimulation. (**C**) Paired comparison of the magnitude of the AIM^+^ Spike-specific CD4^+^ and CD8^+^ T cell responses. (**D**-**E**) Paired comparisons of the magnitude of the Spike-specific T cell responses measured by ICS and AIM assays for (**D**) CD4^+^ T cell and (**E**) CD8^+^ T cell responses. (**F**) Correlation between magnitude of Spike-specific CD4^+^ T cell responses measured by AIM and ICS. (**C**-**F**) include merged data from the two cohorts and both time points. Statistical comparisons were made in **C-E** by Wilcoxon paired tests. In **F**, statistical comparison was made by Spearman test.

**Supplemental Figure 4.**
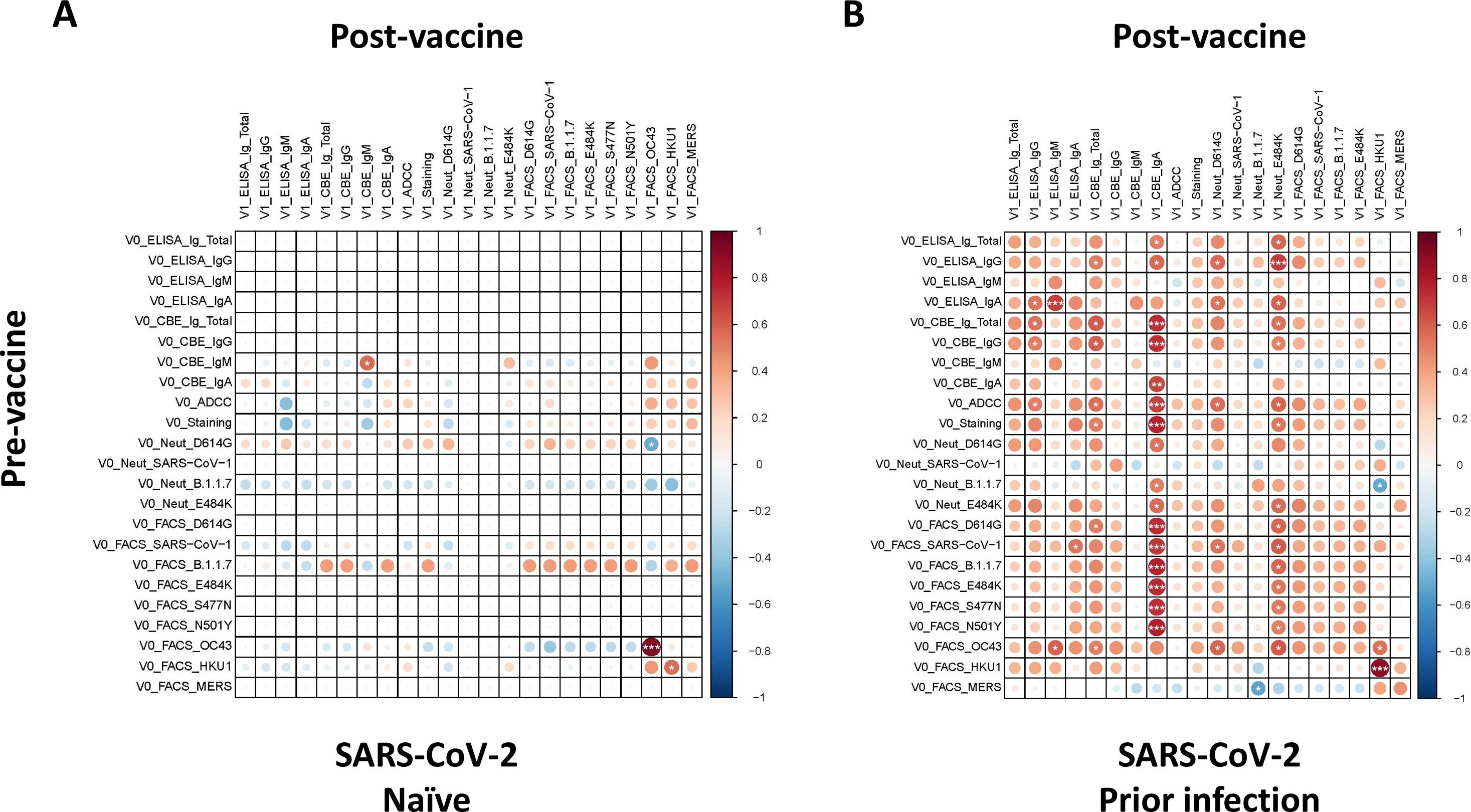
Correlations between serological measurements for induced vaccine responses. Summary of pairwise correlations of humoral parameters between the time point before vaccination against the same responses three weeks post vaccination, both for the naïve (**A**) and prior infection group (**B**). In the correlograms, circles are sized and color-coded according to the magnitude of the correlation coefficient (r). The color code of r values is shown to the right (red colors represent positive, blue colors negative correlations between two parameters). Asterisks indicate statistically significant correlations (*P < 0.05, **P < 0.01, ***P < 0.005). Correlation analysis was done using nonparametric Spearman rank tests.

**Supplemental Figure 5.**
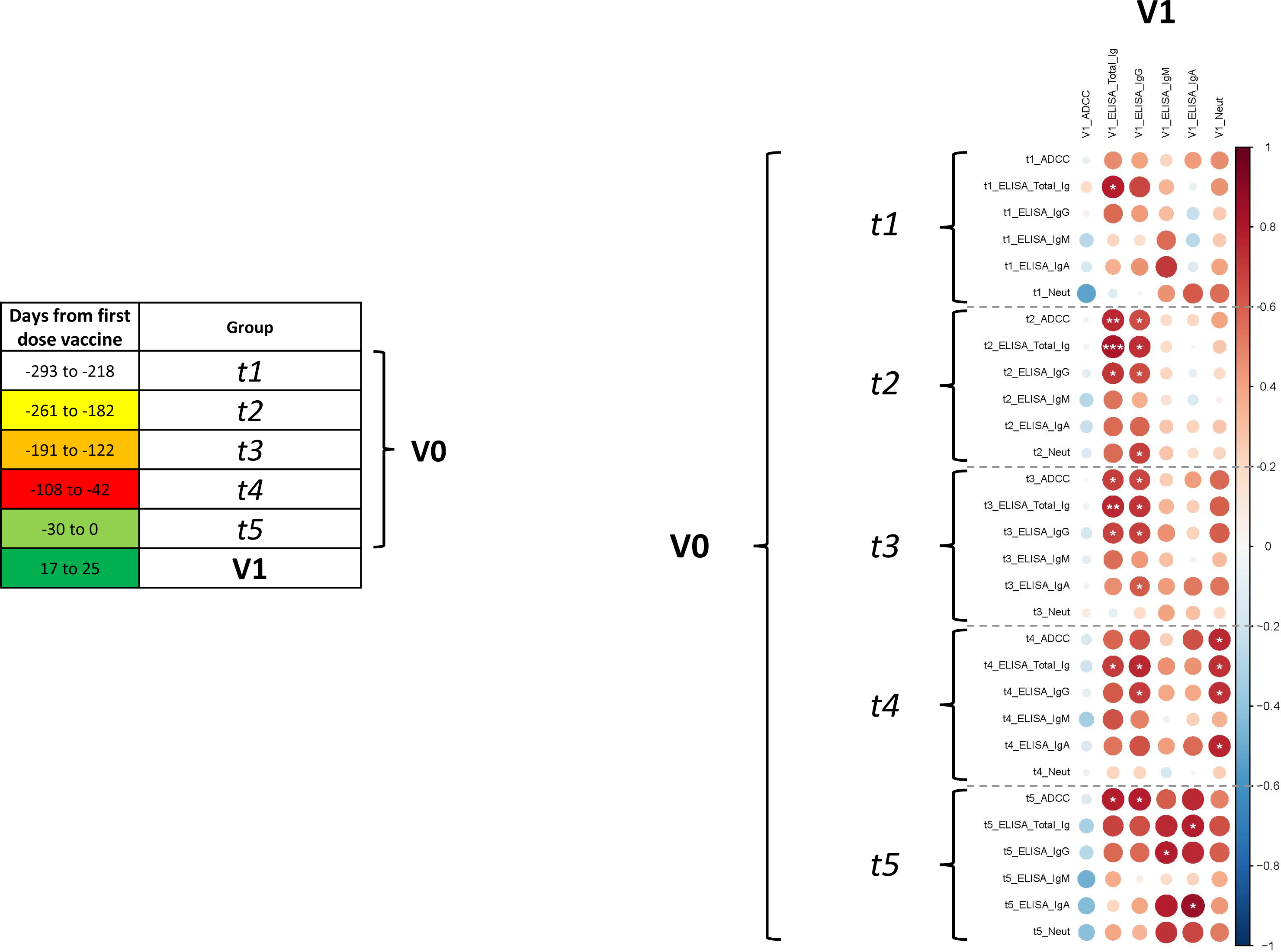
Correlations between longitudinal serological measurements for induced vaccine responses in previously infected SARS-CoV-2 individuals. Summary of pairwise correlations of humoral parameters between longitudinal time points after natural SARS-CoV-2 infection and before vaccination against the same responses post vaccination. In the correlograms, circles are sized and color-coded according to the magnitude of the correlation coefficient (r). The color code of r values is shown to the right (red colors represent positive, blue colors negative correlations between two parameters). Asterisks indicate statistically significant correlations (*P < 0.05, **P < 0.01, ***P < 0.005). Correlation analysis was done using nonparametric Spearman rank tests. Details about the studied time points are provided in the legend to the left.

**Supplemental Table 1.** **Humoral responses before and three weeks after vaccination (Mean +/- SD are shown)**

|  | SARS-CoV-2 Naïve |  | SARS-CoV-2 Prior infection |  |
| --- | --- | --- | --- | --- |
|  | Before Vaccination | After Vaccination | Before Vaccination | After Vaccination |
| **anti-RBD total Ig<sup>a</sup>** | 2.130 ± 0 | 29.823 ± 19.917 | 21.888 ± 17.687 | 113.122 ± 28.754 |
| **anti-RBD IgM<sup>a</sup>** | 3.119 ± 0 | 4.634 ± 4.668 | 4.404 ± 3.385 | 10.331 ± 14.868 |
| **anti-RBD IgA<sup>a</sup>** | 1.227 ± 0 | 2.177 ± 1.644 | 2.103 ± 1.947 | 23.459 ± 16.592 |
| **anti-RBD IgG<sup>a</sup>** | 3.470 ± 0 | 72.180 ± 49.139 | 47.966 ± 38.090 | 236.156 ± 32.851 |
| **anti-Spike total Ig<sup>a</sup>** | 0.043 ± 0 | 1.177 ± 0.787 | 0.974 ± 0.626 | 4.104 ± 0.773 |
| **anti-Spike IgM<sup>a</sup>** | 0.044 ± 0.028 | 0.379 ± 0.666 | 0.089 ± 0.077 | 0.088 ± 0.081 |
| **anti-Spike IgA<sup>a</sup>** | 0.050 ± 0.015 | 0.359 ± 0.356 | 0.287 ± 0.305 | 3.027 ± 2.051 |
| **anti-Spike IgG<sup>a</sup>** | 0.132 ± 0 | 2.811 ± 1.767 | 2.262 ± 1.415 | 9.288 ± 1.777 |
| **Neutralization (D614G) (ID50)** | 53.744 ± 12.823 | 52.495 ± 9.728 | 156.235 ± 172.244 | 2416.848 ± 1752.593 |
| **ADCC (%)** | 2.451 ± 1.176 | 28.355 ± 12.307 | 23.323 ± 9.699 | 66.629 ± 6.161 |

<sup>a</sup>RLU normalized to CR3022, as described in the material and methods section.

**Supplemental Table 2.** **Flow cytometry antibody staining panel for intracellular detection**

| **Target** | **Fluorochrome** | **Clone** | **Supplier** | **Detection** |
| --- | --- | --- | --- | --- |
| **CD3** | BUV395 | UCHT1 | BD | Surface |
| **CD4** | BV711 | L200 | BD | Surface |
| **CD8** | BV570 | RPA-T8 | Biolegend | Surface |
| **CD14** | BUV805 | M5E2 | BD | Surface |
| **CD16** | BV650 | 3G8 | Biolegend | Surface |
| **CD19** | APC-eFluor780 | HIB19 | Thermo Fisher Scientific | Surface |
| **CD40L** | BV421 | TRQP1 | BD | Surface |
| **CD56** | BUV737 | NCAM16.2 | BD | Surface |
| **CD69** | PerCP-eFluor710 | FN50 | Thermo Fisher Scientific | Surface |
| **CD107A** | BV786 | H4A3 | BD | Staining during culture |
| **Granzym B** | Alexa Fluor 700 | GB11 | BD | Intracellular |
| **IFN-g** | PECy7 | B27 | BD | Intracellular |
| **IL-2** | PE-Dazzle 594 | MQ1-17H12 | Biolegend | Intracellular |
| **IL-10** | PE | JES3-9D7 | BD | Intracellular |
| **IL-17A** | eFluor660 | eBio64CAP17 | Thermo Fisher Scientific | Intracellular |
| **TNF-α** | Alexa Fluor 488 | Mab11 | Thermo Fisher Scientific | Intracellular |

**Supplemental Table 3.**
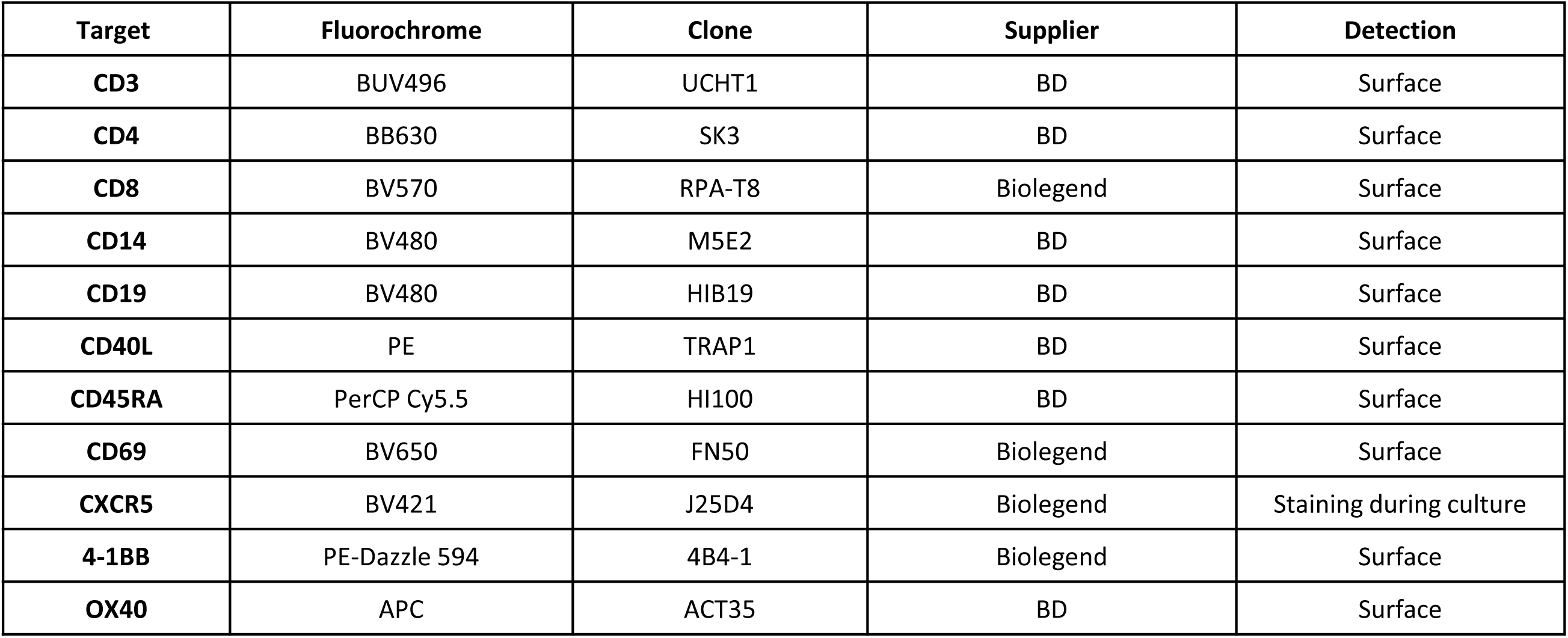
**Flow cytometry antibody staining panel for activation-induced marker assay**

| **Target** | **Fluorochrome** | **Clone** | **Supplier** | **Detection** |
| --- | --- | --- | --- | --- |
| **CD3** | BUV496 | UCHT1 | BD | Surface |
| **CD4** | BB630 | SK3 | BD | Surface |
| **CD8** | BV570 | RPA-T8 | Biolegend | Surface |
| **CD14** | BV480 | M5E2 | BD | Surface |
| **CD19** | BV480 | HIB19 | BD | Surface |
| **CD40L** | PE | TRAP1 | BD | Surface |
| **CD45RA** | PerCP Cy5.5 | HI100 | BD | Surface |
| **CD69** | BV650 | FN50 | Biolegend | Surface |
| **CXCR5** | BV421 | J25D4 | Biolegend | Staining during culture |
| **4-1BB** | PE-Dazzle 594 | 4B4-1 | Biolegend | Surface |
| **OX40** | APC | ACT35 | BD | Surface |

